# “Tumor Treating Fields” delivered via electromagnetic induction have varied effects across glioma cell lines and electric field amplitudes

**DOI:** 10.1101/2023.01.18.524504

**Authors:** Rea Ravin, Teddy X. Cai, Aiguo Li, Nicole Briceno, Randall H. Pursley, Marcial Garmendia-Cedillos, Tom Pohida, Herui Wang, Zhengping Zhuang, Jing Cui, Nicole Y. Morgan, Nathan H. Williamson, Mark R. Gilbert, Peter J. Basser

## Abstract

Previous studies reported that alternating electric fields (EFs) in the intermediate frequency (100 – 300 kHz) and low intensity (1 – 3 V/cm) regime — termed “Tumor Treating Fields” (TTFields) — have a specific, anti-proliferative effect on glioblastoma multiforme (GBM) cells. However, the mechanism(s) of action remain(s) incompletely understood, hindering the clinical adoption of treatments based on TTFields. To advance the study of such treatment *in vitro*, we developed an inductive device to deliver EFs to cell cultures which improves thermal and osmolar regulation compared to prior devices. Using this inductive device, we applied continuous, 200 kHz electromagnetic fields (EMFs) with a radial EF amplitude profile spanning 0 – 6.5 V/cm to cultures of primary rat astrocytes and several human GBM cell lines — U87, U118, GSC827, and GSC923 — for a duration of 72 hours. Cell density was assessed via segmented pixel densities from GFP expression (U87, U118) or from staining (astrocytes, GSC827, GSC923). Further RNA-Seq analyses were performed on GSC827 and GSC923 cells. Treated cultures of all cell lines exhibited little to no change in proliferation at lower EF amplitudes (0 – 3 V/cm). At higher amplitudes (> 4 V/cm), different effects were observed. Apparent cell densities increased (U87), decreased (GSC827, GSC923), or showed little change (U118, astrocytes). RNA-Seq analyses on treated and untreated GSC827 and GSC923 cells revealed differentially expressed gene sets of interest, such as those related to cell cycle control. Up- and down-regulation, however, was not consistent across cell lines nor EF amplitudes. Our results indicate no consistent, anti-proliferative effect of 200 kHz EMFs across GBM cell lines and thus contradict previous *in vitro* findings. Rather, effects varied across different cell lines and EF amplitude regimes, highlighting the need to assess the effect(s) of TTFields and similar treatments on a per cell line basis.

## Introduction

Glioblastoma Multiforme (GBM) is the most common and lethal adult primary brain tumor. Despite a multi-modal treatment plan, patients’ median survival is only 14 – 16 months from the time of diagnosis — a small percentage survive five years or more [1, 12, 41].

Kirson *et al*. [20] reported that intermediate frequency (100 – 300 kHz), low intensity (1 – 3 V/cm) electric fields — termed “Tumor Treating Fields” (TTFields) — have an anti-proliferative effect which inhibits the growth rate of GBM cells in culture while having no noticeable effect on non-dividing cells. This inhibitory effect was reported to be frequency-dependent with a peak at 200 kHz for GBM cells. The effect was also dependent on the electric field (EF) amplitude; at higher EF amplitudes (> 1 V/cm), more than 50% growth rate inhibition was observed in some glioma cell lines [18, 20]. In follow-up studies, TTFields were found to result in decreased animal and human tumorigenesis [11, 18, 20] and metastatic spreading [19]. Abnormal mitotic cell morphology, including blebbing followed by apoptosis, has also been observed under the influence of TTFields [7, 10].

Novocure Ltd. has since translated these preclinical results. The Optune™ system (previously NovoTTF-100A) was developed to apply TTFields *in vivo*. The system consists of an array of electrodes attached to the shaved scalp of the patient. In 2011, the Optune™ system was approved by the U.S. Food and Drug Administration (FDA) for the treatment of recurrent GBM based on a Phase III clinical trial (NCT00379470) comparing TTFields to physician’s choice chemotherapy [43]. In 2015, the Optune™ system with adjuvant temozolomide (TMZ) was approved for the treatment of newly diagnosed GBM following surgery and radiotherapy based on a trial (NCT00916409) comparing TTFields in combination with TMZ to TMZ alone [42].

### TTFields lack a clear mechanism of action

Despite continued clinical, pre-clinical, and theoretical research, the mechanism(s) of action of TTFields remain incompletely understood. Computational models predict that the attenuation of EFs within cells is frequency-dependent [45–47]. At lower frequencies, EFs are “screened” (i.e., impeded by electrical resistance and capacitive reactance) and do not penetrate the cell membrane, whereas in the intermediate (100 – 500 kHz) frequency range, EFs can penetrate with comparatively little amplitude attenuation at different stages of the cell cycle [46]. As a result, TTFields may, in principle, have direct biophysical effect(s) on the interior of GBM cells. Kirson *et al*. [20] initially proposed two effects that were deemed specifically anti-mitotic: (1) TTFields may affect the dipole alignment of charged molecules, namely tubulin and septin dimers, and (2) TTFields may result in non-uniform EFs (i.e., EF gradients) in the cleavage furrow of the dividing cell. The first mechanism would disrupt mitotic spindle formation and possibly the localization of the septin complex to the anaphase spindle midline. The second mechanism would produce significant dielectrophoretic (DEP) forces during cytokinesis, which, in turn, could result in the aggregation of polarizable macromolecules towards and away from the cleavage furrow, disrupting the cleavage process.

Recent computational and theoretical analyses by Tuszyński *et al*. [45] and Li *et al*. [23], however, suggest that clinically relevant EF amplitudes (1 – 3 V/cm) produce electrostatic forces that are too small to significantly affect the rotational dynamics of tubulin or septin — i.e., these forces are much smaller than the thermal energy of said dimers — casting doubt on the first proposed mechanism. As for the second mechanism, these models [45, 46] predict that TTFields can produce significant DEP forces, providing theoretical evidence for the disruptive motion of polarizable macromolecules during cytokinesis. Li *et al*. [23], however, predict that this motion would be too slow compared to the duration of telophase to affect cell division. TTFields may also produce biophysical effects beyond those suggested by Kirson *et al*. [20], including but not limited to: propagating ionic waves along and around microtubule filaments [45], changes in the conductance of ion channels [28], changes in the membrane potential [23], increased membrane permeability [4], and Joule heating of the cytoplasm and/or culture medium [2].

A growing body of work implicates specific biomolecular pathways in the mechanism(s) of action of TTFields (see [37] for review). For example, TTFields applied *in vitro* have been found to result in delayed DNA damage repair via downregulation of BRCA [9], in DNA double-strand breaks [15], and in the enhancement of autophagy via downregulation of the phosphoinositide 3-kinase/Akt/nuclear factor-*κ*B signalling pathway [16]. Other reported effects include enhanced immunogenic cell death [19] and reduced cell migration and invasion [17, 39]. These pathways may operate downstream of — or in tandem with — the aforementioned biophysical effects. TTFields may also have a synergistic effect with adjuvant chemotherapies. The clinical finding that TTFields are most effective in combination with TMZ [42, 43] support this hypothesis. TTFields may enhance the effects of TMZ by interfering with the multi-drug resistance of GBM cells, a feature which is conferred, in part, by the over-expression of ATP-Binding Cassette (ABC) transporters, ABCG2 and ABCB1 [49]. Recent work has also assessed genome-wide expression changes due to TTFields, with effects noted on cell cycle and cell death related genes, among others [3, 22]. While many mechanisms may play a role in the observed effects of TTFields, none has been conclusively demonstrated to operate *in vivo*.

There is an ongoing debate about the efficacy of TTFields and Optune™ within the neuro-oncology community. As of 2019, adoption rates remain around 30% (i.e., 30% of eligible GBM patients receive TTFields where it is available) [37]. This slow adoption might be due to the lack of a clear mechanism of action, discussed above, and/or the absence of a shammed control group in the open-label clinical studies, among other factors (see Wick [48] for a more thorough clinical perspective). To help resolve this debate, Novocure Ltd. has made their *in vitro* test system, called the Inovitro™, available to researchers. Independent studies using the Inovitro™ system to explore various mechanisms of action have since been published (e.g., [15, 32]). As a crucial supplement to these studies, we sought to investigate the effects of intermediate frequency, low amplitude electric fields (EFs) on human GBM cell lines using an altogether different EF application method.

### Redesigning the *in vitro* TTFields experiments

The Inovitro™ system, described in detail in Porat *et al*. [33], is not without methodological challenges. The system consists of two pairs of orthogonally-oriented, capacitively-coupled electrodes attached to cell culture dishes. These electrodes are insulated by ceramic and connected to a function generator and amplifier. Alternating EFs are generated with periodic switching between the electrode pairs [18, 33] to maximize the efficacy of any orientationally-dependent effects, such as DEP forces during cytokinesis.

The major pitfall of the Inovitro™ system is heat generation. A substantial amount of heat is produced via the contacting electrodes and via Joule heating of the (electrically) conductive culture medium. To maintain a physiological temperature of 37 ° C within the culture medium, the developers of the Inovitro™ system suggest that it be operated within an 18 °C incubator. Furthermore, the system is designed to be intermittently switched off [33]. The use of a refrigerated environment introduces additional challenges. The cooler, drier air surrounding the warm culture medium may accelerate evaporation due to increased vapor pressure gradients. As a result, osmolar regulation may be compromised over longer periods of time.

In developing our EF applicator device, we considered the relevant design limitations of the Inovitro™ system. To mitigate the difficulties associated with heat generation, we chose to use an inductive — and thus non-contacting — EF delivery method. In this way, conductive heating is eliminated. The setup is detailed in Ravin, Cai *et al*. [35]. In brief, electromagnetic fields (EMFs) at the desired frequency (200 kHz) were generated within an air-core solenoid coil connected to an industrial induction heater and placed within an incubator (see Methods, Fig 1A). An induced, circumferential EF profile is produced, with an amplitude that increases linearly from the center of the coil (Fig 1B). The radially increasing EF profile within each culture dish enables us to perform a controlled EF “dose titration” experiment, wherein each dish has a core region of little to no EFs that serves as an internal control. Any between-dish confounds, such as differences in osmolar conditions, cannot contribute to differences observed within the same dish, and were thus avoided. A small, vertically (or axially) alternating magnetic field is also produced. Joule heating of the culture medium, however, remains an issue.

**Figure 1.**
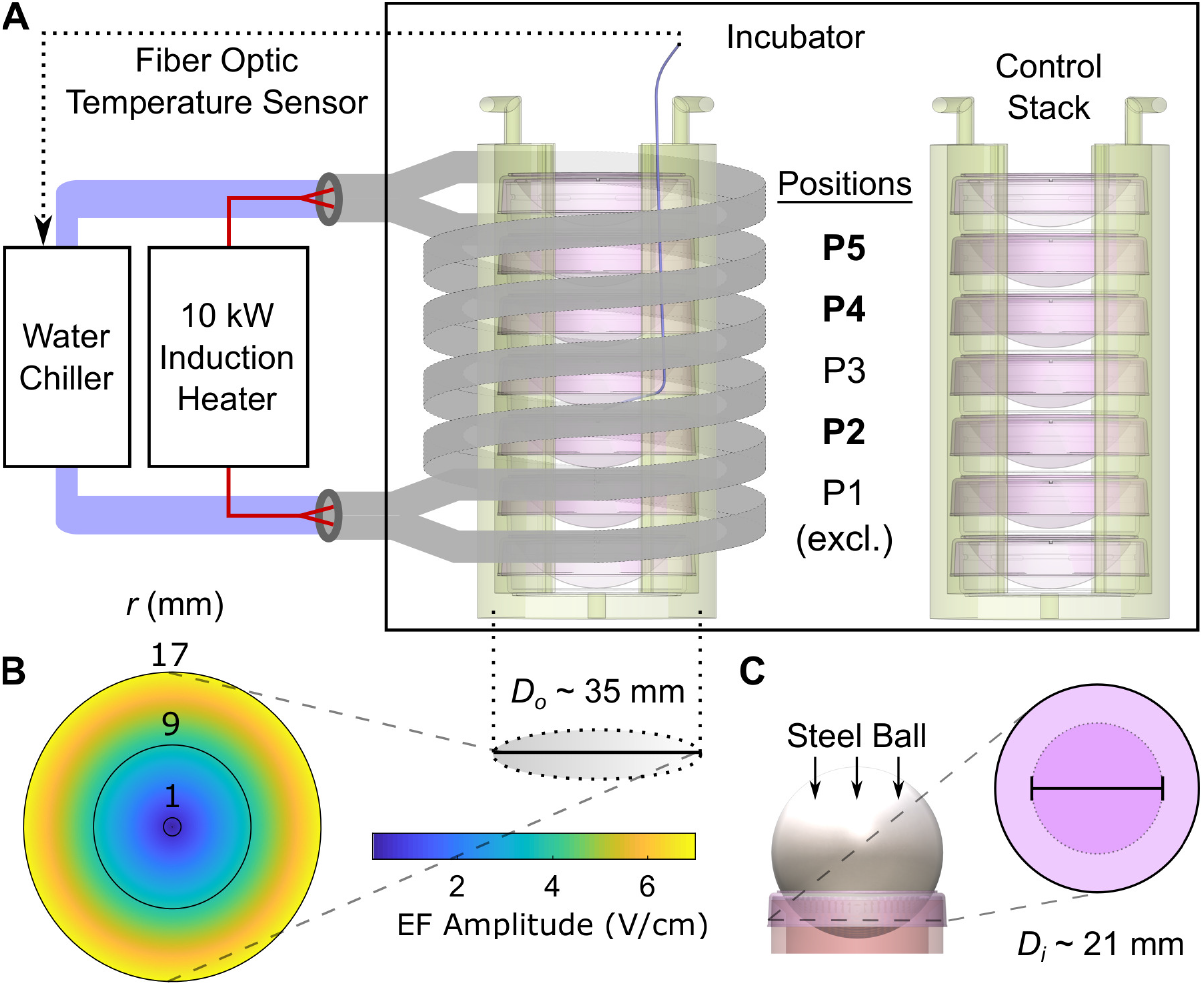
Schematic of experimental apparatus and methods. (A) Setup for application of 200 kHz EMFs. The copper coil (inner diameter = 5 cm, height = 8 cm, 3 turns, double-wrapped) was attached to an induction heater and water-jacketed. Water was circulated using a chiller loop. The coil was then fixed inside an incubator. Sleeves with seven stacked dishes were placed within the coil, five of which were situated within the coil at vertical positions labelled P1 – P5. P1 was excluded due to observed temperature variability. Temperature was monitored using a fiber optic sensor in the center of the P3 dish; P3 was thus excluded as well. A separate control stack was placed away from the coil. (B) Measured EF amplitude profile. Outer diameter *D_o_* = 35 mm is overlaid. (C) Diagram of lid shaping process used to reduce the exposed surface area. The lids were shaped to contact an inner diameter of approximately *D_i_* = 21 mm.

To address Joule heating, the coil was actively cooled below the temperature 120 of the incubator, thus producing a net convective heat loss that can be adjusted 121 to balance the Joule heating of the culture medium and maintain a steady-state 122 temperature. In doing so, EMFs can be applied continuously without affecting the temperature of the culture medium. Steps were also taken to improve osmolar regulation (see Methods, Fig 1C).

This experimental test system allows us to evaluate the results reported by Kirson *et al*. [20] as well as extend the EF “dose” or amplitude range beyond those previously used. Here, we report the results of 72 hours of continuous 200 kHz EMF application spanning 0 – 6.5 V/cm on various cell lines. To our knowledge, we report the first results of alternating EF stimulation in excess of 5 V/cm. We studied two human GBM cell lines utilized in prior studies: U87 and U118, and two recently validated GBM stem cell lines: GSC827, GSC923 [36]. Primary rat astrocytes were also studied as an example of normal central nervous system cells. Segmented pixel density was assessed via GFP expression for U87 and U118 and via CellTracker Green CMFDA and Hoechst 33342 staining for GSC827, GSC923, and rat astrocytes as a proximal measure of cell density. The time-course of CellTracker and Hoechst staining intensity in GSC827 and GSC923 cells was also assessed to investigate whether the treatment affects the activity of pumps that remove CellTracker and Hoechst, namely ABC transporters (ABC transporter activity has been postulated as a source of multi-drug resistance in GBM [49]). Finally, RNA-Seq analyses were performed on GSC827 and GSC923 cells drawn from different radial dish regions — corresponding to different EF amplitudes — to assess effects on gene expression following treatment.

## Materials and Methods

### Cell cultures

The U87 and U118 cells expressed GFP, whereas GSC827, GSC923 and rat astrocyte cells did not. For all cultures, 3 mL of cell suspension were placed in 35 mm culture dishes (Corning, Falcon). For U87, U118, GSC827, GSC923, and rat astrocytes, the plating density in cells per dish was 1.0 × 10^5^, 5 × 10^4^, 7.5 × 10^4^, 7.5 × 10^4^, and 1.0 × 10^5^, respectively. The number of replicates was 21, 24, 18, 12, and 15, in the same order.

The U87 and U118 cells (ATCC, Glioma Tumor Cell Panel TCP-1018™) were maintained in 75 ml flasks with DMEM culture media supplemented with 10% FBS, 1% penicillin-streptomycin (Gibco; Thermo Fisher Scientific). Cells were passaged twice a week and cell cultures were maintained in a 5% CO_2_ incubator at 37 °C. Before each experiment, GFP-labeled GBM cells were detached from the culture vessel with Accutase™ (Sigma) and re-suspended in media. To aid in cell attachment, cells were plated on dishes coated with Poly-D-Lysine (0.05 mg/ml) (Sigma).

To elucidate the effects of 200 kHz induced EMFs on brain tumor transcriptomic profiles, we performed RNA-Seq on cultures of the GSC827 and GSC923 cell lines, which were originated from the Neuro-Oncology Branch [36], rather than traditional model cell lines; GSC827 was derived from a 60-year-old male GBM patient and GSC923 was from a 56-year-old female GBM patient. These cell lines better represent the heterogeneity that may be found amongst GBM patients. The GSC827 and GSC923 cells were grown in suspension in 75 ml flasks with NBE medium composed of DMEM/F12 medium supplemented with 10 ml P/S, 5 mL L-Glutamine, 5 ml N2 Growth Supplement, 10 ml B27 Growth Supplement, EGF (25 ng/ml), and bFGF (25 ng/ml). Cells were passaged once a week, dissociating the spheres that form with Accutase™. Between each passage, media was changed once. Cells were plated on dishes coated with Matrigel (0.4 mg/ml, 356234 Corning), again to aid in attachment.

Astrocytes were isolated from the cortices of postnatal rats between 1 and 3 days after birth. Astrocytes were cultured in DMEM + 10% FBS at 10% CO_2_. After 7 days of growth, flasks were shaken overnight to reduce contamination by oligodendrocyte progenitor cells and microglia. Once the cultures had reached 75% confluence (approximately 3 weeks), cells were lifted with Accutase™ and then frozen. Before experiments, cells were thawed and plated in a 75 ml flask with DMEM. At 75% confluence, cells were lifted with Accutase™ and plated on dishes coated with Poly-D-Lysine.

### Continuous delivery of 200 kHz EMFs to cell cultures

The test system, schematized in Fig 1, is described in detail in Ravin, Cai *et al*. [35]. In brief, the system was powered by an industrial 10 kW induction heater (DP-10-400, RDO Induction LLC, Washington NJ) based on a conventional LC resonant circuit design. The induction device was connected to an air-core double copper coil. The coil was jacketed and connected to a water chiller (Durachill©, PolyScience) to remove heat and thereby control the coil temperature (Fig 1A). The coil was fixed inside of a 5% CO_2_ incubator (MCO-18M Multigas Incubator, SANYO Electric Co. Ltd., Osaka, Japan) through a Plexiglas window fixture. Plastic sleeves for cell culture dishes were 3-D printed in-house. Each sleeve accommodated seven 35 mm dishes in a vertical stack. The top and bottom plates at the edges of the coil were filled with only media; the middle five plates contained cell cultures and their positions were labelled P1 – P5 (Fig 1A).

The coil was actively water-cooled to a temperature of about 22 °C, subject to adjustment, in order to remove the heat generated by Joule heating via steady-state convection and thus maintain a constant, physiological temperature within the culture media. The experimental temperature was measured continuously using a fiber optic temperature sensor placed in the center of the dish at P3 within the coil, which contained only culture medium (Fig 1A). This temperature was remotely monitored, controlled, and logged. Stable temperature conditions (37 ± 0.5 °C) were maintained during all experiments. Similar temperatures were maintained across the dishes at positions P2 – P5, but not at P1, which was measured to be cooler by several °C [35]. Dishes at P1 were excluded from all analyses. Each 72 hour experiment produced three experimental replicates of pair-wise dish comparisons at positions: P2, P4, P5.

The expected EF profile generated within the coil by the induction device can be calculated from basic principles of electricity and magnetism. Inside the coil, the induced EF amplitude increases linearly from zero at the center of the dish/coil, *r* = 0, to a maximum at *r* = *R*, the outer radius of the dish. While the EF amplitude varies spatially in the radial direction, it alternates in the azimuthal or circumferential direction. At the utilized power level of 3.19 kW, we measured a maximum EF amplitude of 6 – 7 V/cm pk-pk at the dish periphery, illustrated in Fig 1B. This EF profile was measured experimentally using wire loops [35] and a linear fit was obtained for the EF amplitude, |**E**|, as a function of *r* (see Fig 1B),

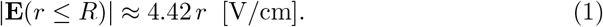

The control sleeve was placed in the same incubator far from the coil and off of its vertical axis (see Fig 1 in [35]) to avoid extant or stray EMFs.

No media changes were performed during experiments. To mitigate media loss, the surface-to-volume ratio of the media was reduced by two methods: (1) by increasing the volume of media in the plate from the conventional 2 ml to 3 ml and (2) by re-shaping the lid of all dishes so as to occlude a portion of the exposed surface area and to redirect condensate back into the media (Fig 1C). Re-shaping was performed by heating the lids using a heat gun and then molding them using a steel ball. A decrease in surface area of ≈ 36% was achieved. With these adjustments, osmolarity increases over the course of 72 hours were measured to be 17% and 6.5% from an initial osmolarity of 314 millilosmoles in the treated and control conditions, respectively [35].

### Confocal imaging and staining

The radially-varying EF amplitude precludes the use of conventional cell lifting and counting methods because such methods discard critical spatial information. Instead, entire dishes were imaged *in situ* by stitching patches together. At the end of experiments, dishes were gently removed from the sleeves. Each pair of dishes (control, treated) was imaged alternately. The rotational orientation of each dish was retained using the manufacturer’s dish markings, i.e., all dishes had indexing markings, and were positioned to face in the same relative direction in the incubator and the imaging stage.

Dishes were mounted to the stage of a Zeiss LSM 780 laser scanning confocal microscope system. For imaging, an EC Plan-NeoFluor 5×, 0.16 N.A. was used. The U87 and U118 cultures were imaged for GFP. For the GSC827, GSC923 and rat astrocyte cultures, NBE medium was removed and 1 ml of NBE medium with CellTracker Green CMFDA (C7025, Thermo Fisher) at a concentration of 2.5 *μ*M or with Hoechst 33342 at a concentration of 3 *μ*g/mL was added prior to imaging. For both GFP and CellTracker, a 488 nm laser, a beam splitter of 488/562 nm, and filters from 500 – 555 nm were used. For Hoechst 33342, a 405 nm excitation wavelength and notch reflector of 405 nm were used. For all images, a total of 484 areas of interest were stitched together with 5% overlap. The final image size was 10,726 × 10,726 pixels (px). Care was taken to use non-saturating parameters for imaging; parameters were kept constant through all experiments.

To assess the ability of GSC827 and GSC923 cells to pump out CellTracker and Hoechst, additional imaging studies for both CellTracker and Hoechst retention were performed after some experiments. Cells were washed twice with 1 ml NBE medium after initial imaging and then placed back into the incubator for 2 hours (without treatment). At the end of 2 hours, dishes were imaged again with the same protocol and imaging parameters.

### Image processing

Image processing aimed to quantify segmented px density as a function of radial distance. Each image was first divided (i.e., binned) into 25 equal annular length rings or bands. Due to fitments which occlude the bottom portion of the dish, only an upper portion of the band was included in the analysis from the 18th band onward (see Fig S1 for an exemplar full dish image). Due to deformations at the edge of the dish, the 23rd – 25th bands were also discarded. Images were first contrast enhanced using a morphological transform method in which the “top hat” is added and the “bottom hat” is subtracted from the image. A disk-shaped structuring element with a 3 px radius was used for this step. Images were then denoised using a 3× 3 Wiener filter.

Next, segmentation was performed on a band-by-band basis in two steps: binarization and opening (Fig 2C). Thresholds for image binarization were selected on a band-by-band basis using Otsu’s method. For all cell types except U87, a single threshold was calculated. For U87, two thresholds were calculated and the lower of the two was used because of the presence of brighter cells exhibiting high GFP expression. Following binarization, an opening step was performed with a 6 px radius. Finally, the ratio of foreground and background px was calculated for each band as a proximal measure of the cell density. The foreground pxs were used as a mask for the later calculation of band-by-band px intensities to assess pumping activity. All image processing was performed using MATLAB 2019a (Natick, MA). See Methods in [35] for further details and representative code.

**Figure 2.**
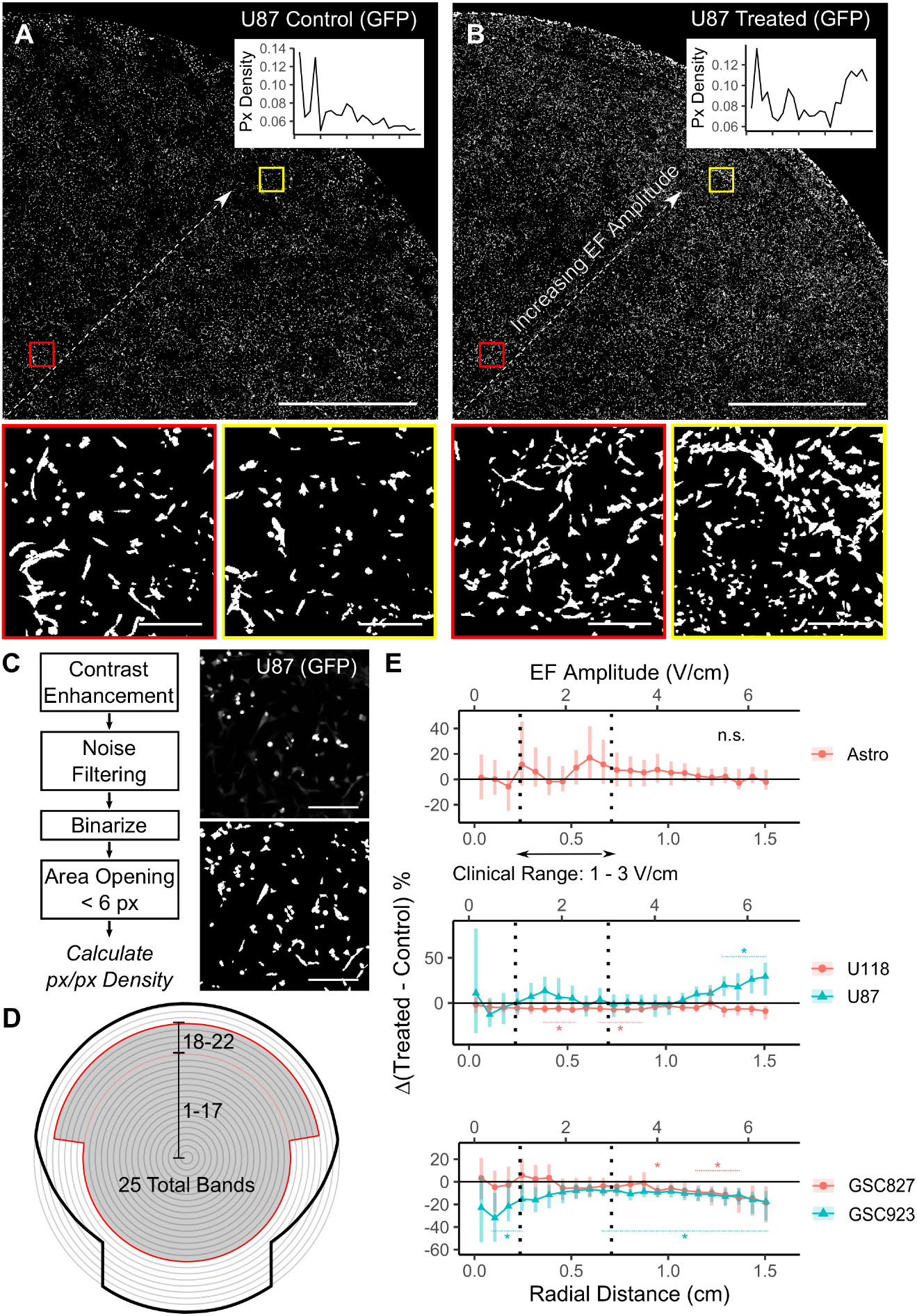
Summary of segmented px density results. (A) Exemplar image of the upper right quadrant of a U87 control dish after all image processing steps (scale bar = 5 mm). Inset graph shows the calculated px densities from bands 1 – 22 (see panel D). Zoomed patches near the center and edge are shown (scale bar = 0.25 mm). (B) Corresponding treated dish. Note the increasing trend in px density with EF amplitude. (C) Image processing steps. To the right, an exemplar raw image for U87 cells (top) is compared to the final, binarized image (bottom). (D) Diagram of the discretization of the dish into 25 equal annular bands. The included area is shaded. (E) Aggregated results. The difference between normalized px densities for different cell lines is plotted as a function of both radial distance (bottom axis) and EF amplitude (top axis) using Eq. (1) for axis conversion. Error bars = 90% confidence intervals from bootstrapped pair-wise differences with 1000 re-samplings. One-sample Wilcoxon signed rank tests were also performed band-wise to find significant changes in the treated vs. control difference (* = *p* < 0.05). The number of replicates was *N* = 15, 24, 21, 12, 18 for, in legend order, rat astrocytes (Astro), U118, U87, GSC827, and GSC923, respectively. Note that the smaller bands are expected to have fewer cells and thus higher variability.

Analyses of the images was performed in a pair-wise manner, i.e., between each pair of control and treated cultures in the same coil position (P2, P4, P5). Px densities were normalized to (i.e., divided by) the mean px density of each control dish across all included bands before pooling results across pairs for further statistical analyses.

To validate a correspondence between px density and experimental cell count, additional control experiments on U87 cells were performed. In these experiments, a range of plating densities was used and cells were lifted and counted after imaging. A linear relationship was observed between the experimental cell count and the total (i.e., without binning) segmented px density (Fig S2). The segmented px density determined in this way thus serves as a robust proxy of cell density (see also Fig 6 in [35]).

### RNA extraction

RNA-Seq analyses were performed on GSC827 and GSC923 cells from 3 experiments (N = 9) each. At the end of these experiments, media was replaced with 1 ml of Accutase™ for several minutes until the cells began to round up. Then, 3 ml of DMEM/F12++ was added to stop the cells from lifting off. Using a 200 *μ*L pipettor, cells were manually collected from 3 regions in each dish: (1) from a radius of 4 mm around the center of the dish, and then from 3 mm bands centered at a radial distance of (2) 8.5 mm and (3) 15.5 mm from the center of the dish. Again, these regions are labelled as the treated center (TC), treated middle (TM), and treated surroundings (TS), respectively, and likewise for control cultures (CC, CM, CS). Regions were kept consistent using a traced diagram.

Cells were spun down and re-suspended in DMEM/F12++ and kept on ice. Samples were centrifuged at 800 ×*g* for 5 minutes at 18°C. Media was aspirated and the cell pellet was re-suspended in 350 *μ*L RLT buffer (Qiagen RNeasy Mini Kit, No. 74104). Extraction was automated using a QIAcube set to the RNA-RNeasy mini protocol with DNaseI digest and 35 *μ*L DEPC water elution. RNA quantity was measured using Qubit RNA broad range kit (Thermo Fisher, No. Q10211) using 1 *μ*L of the sample. Quality was determined by running 1 *μ*L of the sample on a BioAnalyzer RNA Nano chip (Agilent, No. 5067-1511) for RIN. The transcriptomic library was prepared using the Illumina TrueSeq stranded mRNA library Prep kit following the manufacturer’s instructions. The library was sequenced on an Illumina NovaSeq SP platform using 150 bp paired-end reads carried out by NCI’s Advanced Technology Research Facilities.

### RNA-seq analysis and identification of differentially-expressed genes

Strand-specific RNA-seq reads were analyzed using CCBR Pipeliner (github.com/CCBR/Pipeliner). This pipeline performs several tasks: pre-alignment reads quality control, grooming of sequencing reads, alignment to human reference genome, post-alignment reads quality control, feature quantification, and differentially-expressed gene (DEG) identification. In the quality control phase, the sequencing quality of each sample was independently assessed using FastQC, Preseq, Picard tools, RSeQC, SAMtools and QualiMap. The samples had 44 to 55 million passed-filter reads with more than 92% bases above the quality score of Q30. Sequencing reads that passed quality control thresholds were trimmed for adaptor sequences using the Cutadapt algorithm with the reference genome (Human - hg38). The transcripts were annotated and quantified using STAR. DEGs were determined using the EdgeR, DESeq2, and limma/voom methods. We derived DEGs using thresholds of *p* ≤ 0.05 and fold changes ≥ 1.3 or ≤ −1.3 across all three methods by comparing and contrasting the regions TS vs. TC, TM vs. TC, and TS vs. TM. Since the number of DEGs was small, the DEGs derived from EdgeR using the same thresholds were also used for functional analysis. We also derived DEGs by contrasting TS vs. CS, TM vs. CM and TC vs. CC using the same thresholds as other contrasts. Genes showing expression levels that were correlated with the EF amplitude region (i.e., TC < TM < TS or TC > TM > TS) were identified using a Pearson correlation with a threshold of *p* < 0.05 and a correlation coefficient of *r* > 0.45.

### Pathway and network analysis

Pathway and network analysis was carried out using Ingenuity Pathway Analysis (IPA; qiagenbioinformatics.com), and Cytoscape (cytoscape.org). More than 2,000 DEGs were uploaded into IPA and core analysis was performed for each contrast using both up-and down-regulated genes. The outputs from canonical pathways, upstream analysis, diseases and functions, and regulator analysis were inspected to elucidate potential underlying molecular functions. In addition, pathway overlay, and network analysis was used to further expand our understanding of biological function. For functional analysis in Cytoscape, the GBM network was downloaded from the STRING database and genes correlated with the EF amplitude region were overlaid for building the corresponding networks.

### Gene set enrichment analysis (GSEA)

We carried out GSEA by contrasting TS vs. TC, TM vs. TC, TS vs. TM, TS vs. CS, TM vs. CM and TC vs. CC. (http://software.broadinstitute.org/gsea/index.jsp). GSEA analysis was performed using the JavaGSEA desktop application (GSEA ver. 4.1). For filtering GSEA databases, MSigDB C2 and C5 curated gene sets of less than 10 genes or greater than 500 genes were excluded. The *p*-values were calculated by permuting the samples 1,000 times to identify enriched gene sets. A nominal *p*-value of less than 0.05 and a normalized enrichment score (ES) greater than 1.4 from GSEA outputs was used as a threshold to determine up- or down-regulated gene sets for each comparison or contrast. R (ver. 4.0.5) was used to cluster and summarize the GSEA outputs.

## Results

### Treatment with 200 kHz induced EMFs produce varied effects on the growth rate of different cell lines

Results are represented as a percentage difference between the final treated vs. control segmented pixel densities, expressed as a function of the radial band, or, equivalently, the EF amplitude (Fig 1B). An example pair of control and treated U87 cultures is shown in Figs 2A – B after all image processing steps. The displayed images provide an example of a radially-dependent cell density increase in the treated U87 dish, especially for EF amplitudes exceeding 4 V/cm. See Methods for details on image processing and segmentation steps (Figs 2C – D). Effects at different EF amplitudes were found to vary for the different cell lines (Fig 2E). Rat astrocytes and U118 cells show little to no change at any EF amplitude, although U118 shows a small (< 5%) difference throughout which is significant in some bands; GSC827 and GSC923 cells show a decrease of approximately 10 – 20%, but only at larger EF amplitudes (4 – 6 V/cm); and U87 cells show an increase of up to 30% at larger EF amplitudes. Results for Hoechst staining (U118, GSC827, GSC923, astrocytes) were similar with the exception of GSC827, which showed no change in segmented px density when measured by Hoechst staining. These results are presented in the Supplement (Fig S3).

Note that the results for astrocytes, GSC827, and GSC923 cells represent the response of only live cells because CellTracker is only taken up by live cells. For some experiments with U87 and U118 cells, additional staining was performed with propidium iodide (PI), a dye permeant only to dead cells. In all such experiments, relatively low numbers of dead cells were found, indicating that cell death was not a contributing factor to the reported effects (see Fig S4 for protocol and exemplar results of PI staining). For some experiments with U118 and rat astrocyte cells, additional staining was performed with Annexin V, which conjugates to apoptotic cells. Few apoptotic cells were found in rat astrocyte cultures. Some apoptotic cells were found in U118 cultures (< 0.5% px/px, or about 3% of the segmented px density from GFP), but no difference was found between treated and control cultures (see Fig S5 for protocol and results of Annexin V staining).

### Apparent pumping activity is only slightly reduced by treatment in GSC827 and GSC923 cells

To evaluate the effect of 200 kHz induced EMFs on pumping activity, the time-course of the intensity of CellTracker and Hoechst staining was studied in GSC827 and GSC923 cultures. CellTracker is pumped by the ABCC1/ABCG2 transporters and poorly by ABCB1 [40]. Hoechst is pumped by ABCG2 and ABCB1 [40].

The reduction in px intensity after 2 hours was assessed for GSC827 and GSC923 cultures using the foreground px for each measurement, as obtained in Fig 2, as a mask. In all culture dishes, a large reduction in intensity (≈ 75%, ~ 0.15 a.u.) was observed for CellTracker, but little to no reduction was observed for Hoechst (< 5%, ≲ 0.01 a.u.) across the dish. An example region is shown in Fig 3A. The difference in CellTracker intensity after 2 hours, Δ, was calculated for each dish on a band-by-band basis. These Δ values were then compared between treated and control dishes as a ratio: Δ_Treated_/Δ_Control_. Results are presented in Fig 3B. For both cell lines, the apparent pumping activity of CellTracker was only slightly reduced in the treated dishes at moderate EF amplitudes (> 2 V/cm). The reduction in intensity for Hoechst was too small to be meaningfully compared in the same way without outlier rejection. Analogous results for Hoechst are presented in Fig S6; no difference in apparent pumping activity was observed.

**Figure 3.**
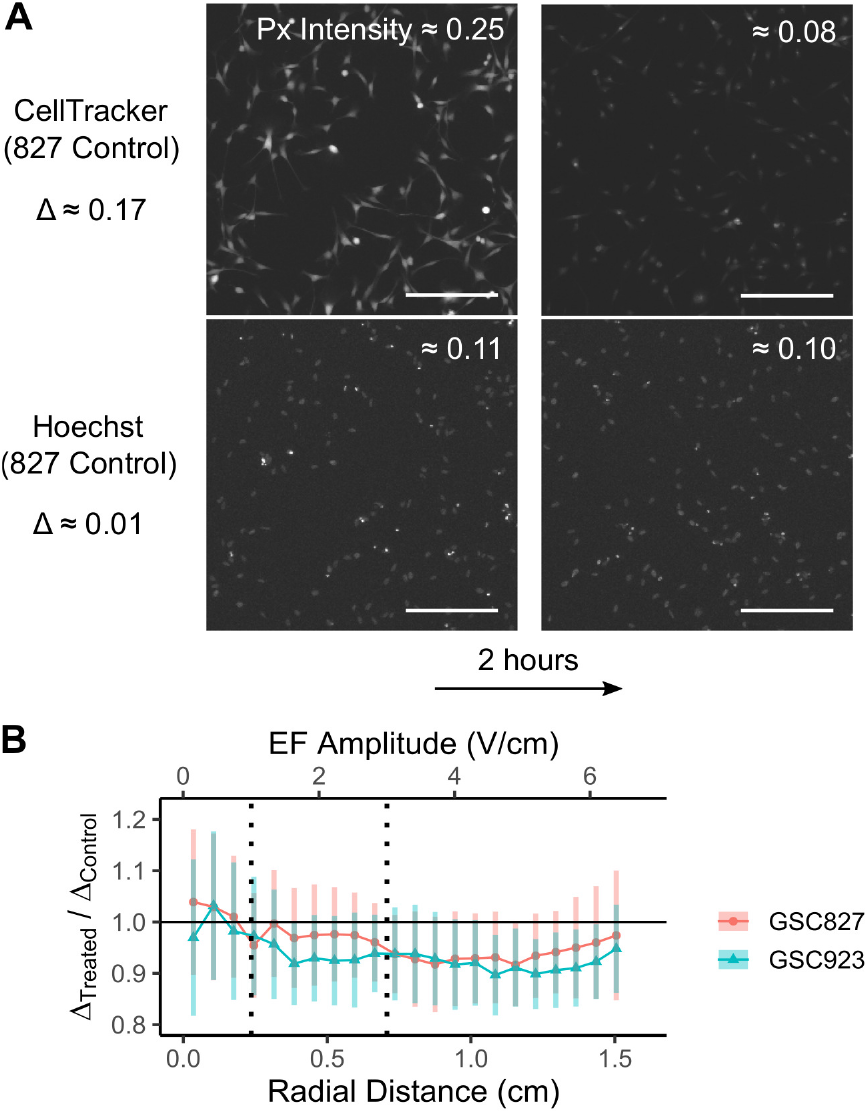
Results for pumping activity. (A) Exemplar images taken from near the center region of a control GSC827 dish (scale bar = 0.25 mm). All images are shown with a flat, normalized brightness increase of 0.15 to aid visualization. Note that the decrease in segmented px intensity, Δ, is much greater for CellTracker than for Hoechst. (B) Ratio of the measured intensity drop of CellTracker over 2 hours, Δ, for treated and control conditions in GSC827 and GSC923. *N* = 12, 18 for GSC827 and GSC923, respectively. Error bars = 90% confidence intervals from bootstrapped pair-wise differences with 1000 re-samplings.

### Transcriptomic differences between the human glioma stem cell lines, GSC827 and GSC923, are greater than the effects of treatment

Analyses were carried out on cells collected from 3 regions in each culture dish: (1) near the center, (2) between the center and the edge, and (3) near the edge. These regions are labelled as the treated center (TC), treated middle (TM), and treated surroundings (TS), respectively (see Methods, Fig 4A). Throughout, these regions are referred to as “EF amplitude regions”. The average EF amplitude 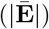 in each region is 0.85, 3.61, and 6.58 V/cm, respectively (Fig 4A). Comparisons were performed between pairs of regions (e.g., TS vs. TC), forming different pairwise contrasts. An identical division of dish regions was used for comparison to control dishes, labelled as control center (CC), control middle (CM), and control surroundings (CS), respectively.

**Figure 4.**
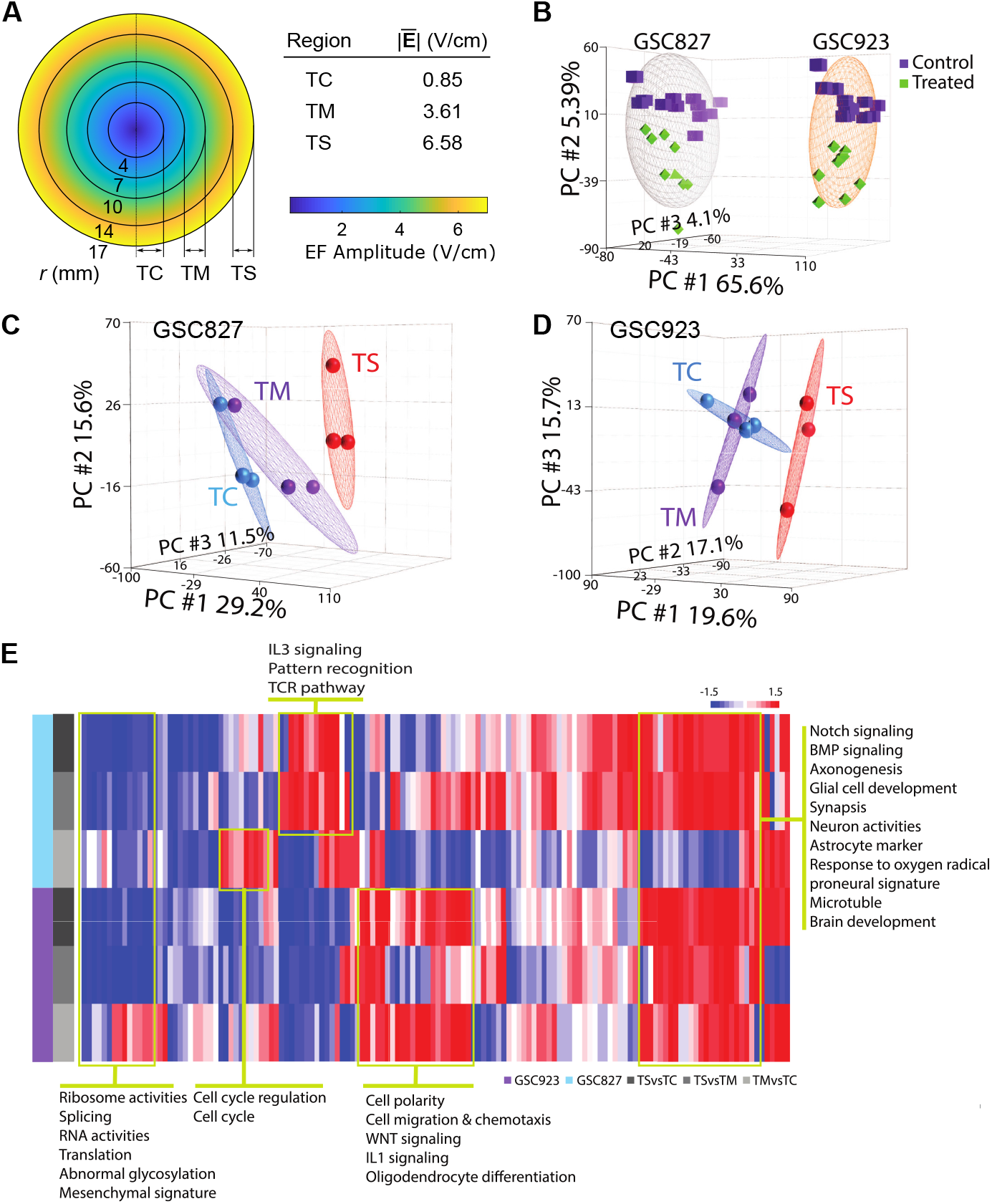
RNA-Seq experimental design and high-level functions affected by different regions of EF amplitude exposure for two glioma stem cell lines, GSC827 and GSC923. (A) Treatment schema and dish arrangement. The treated dish is divided into three discrete regions: TC, TM, and TS. The average EF amplitude in each region 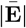 is shown to the right. (B) Unsupervised principal component analysis (PCA) of all samples for GSC827 and GSC923 while pooling treated (TC, TM, TS) and control regions (CC, CM, CS). Ellipses highlight the separation between cell lines. (C–D) Unsupervised PCA of GSC827 and GSC923 separated by the treated dish region. (E) Functional summary of the gene set enrichment analysis for the TS vs. TC, TS vs. TM, and TM vs. TC contrasts in both cell lines.

An unsupervised principal component analysis (PCA) reveals that the samples from these two cell lines are projected into two separated clusters and that treatment effects, which are aligned with the cell lines, are much smaller than the intrinsic differences between the cell lines (Fig 4B–D). The unsupervised PCA indicates that TS is clearly separated from TC and TM, but the global difference between TC and TM in both cell lines is small (Fig 4C–D). Results from the GSEA are summarized in Fig 4E; effects on gene sets of particular interest, such as those related to cell cycle regulation, are labelled.

Note that for all comparisons, the proportion of commonly upregulated or downregulated DEGs between GSC827 and GSC923 is small, ranging from 2.1 – 4.3% of identified genes (Fig S7A). Consistent with these findings, the overlap of enriched gene sets in the TS vs. TC contrast from MSigDB C2 and C5 databases is poor, ranging from 2.1 – 7.6% (Fig S7B). These findings indicate that the two cell lines possess globally different gene expression profiles and are transcriptomically different entities, though both are glioma stem cell lines derived from GBM patients.

### Transcriptomic differences between control cultures and the center of the dish in treated cultures are greater than the effects of treatment

Cells were also cultured in control dishes placed in the same incubator, but far from the coil to avoid EMF exposure. These dishes serve as a separate control (from TC) to evaluate any potential confounding effects, such as diminished osmolarity regulation — an effect quantified in our previous publication [35]. Unsupervised PCA indicates that the gene expression profiles in all three regions of the control dishes (CC, CM, CS) for GSC827 and GSC923 are overlapped but significantly separated from TC (Figs S8A – D). The average EF amplitude in the TC region is 0.85 V/cm (Figs 1A, 4A), which is not expected to be a sufficient EF amplitude to elicit substantive effects on gene expression nor cell growth. Surprisingly, then, the difference in gene expression between the control dishes (CC, CM, CS) and TC is qualitatively larger than the difference between regions in the treated dishes (TC vs. TM, TC vs. TS). This difference suggests that the effect of 200 kHz EMFs on gene expression per se was smaller than the potential effects of osmolarity, the alternating magnetic field, or other experimental differences. That said, we chose to use TC as our reference in all further statistical comparisons to measure the effects of treatment in this study, thus avoiding these potential dish-to-dish confounds in the RNA-Seq analyses. For completeness, we also provide analyses using the control dish as the reference (i.e., TS vs. CS, TM vs. CM, and TC vs. CC); these findings largely overlapped with the findings using TC as the reference (Fig S9).

### Treatment transformed GSC827 and GSC923 cells from expressing mesenchymal to proneural enriched features

Gene set enrichment analysis (GSEA) found that treatment transformed GSC827 and GSC923 cells from expressing mesenchymal enriched features to proneural enriched features (Figs 5A–B). The effects were significant in the TS vs. TC and TS vs. TM contrasts, but not in the TM vs. TC contrast (Fig 5C). There were approximately forty proneural and mesenchymal signature genes significantly correlated with the EF amplitude regions for each cell line (Figs S10A–C), but these genes poorly overlapped between the two cell lines; only 8 genes were shared between the two cell lines (Fig S10B). BCAN, a proneural signature gene, was positively correlated with the EF amplitude regions in both cell lines (Figs 5B, S10D). ITAG5, a prognostic mesenchymal signature gene, was negatively correlated with the EF intensity regions in both cell lines (Figs 5G, S10D). PDPN and PLAUR, which induce mesenchymal gene expression and promote epithelial-mesenchymal transition, were downregulated only in the TS region (Figs 5G, S10D). The treatment also changed the expression pattern from a GBM enriched signature to anaplastic oligodendroglioma (AO) and pilocytic astrocytoma (PA) signatures in GSC827 (Fig 5C). In GSC923, the changes revealed were in the same direction, but were not significant (Fig 5C).

**Figure 5.**
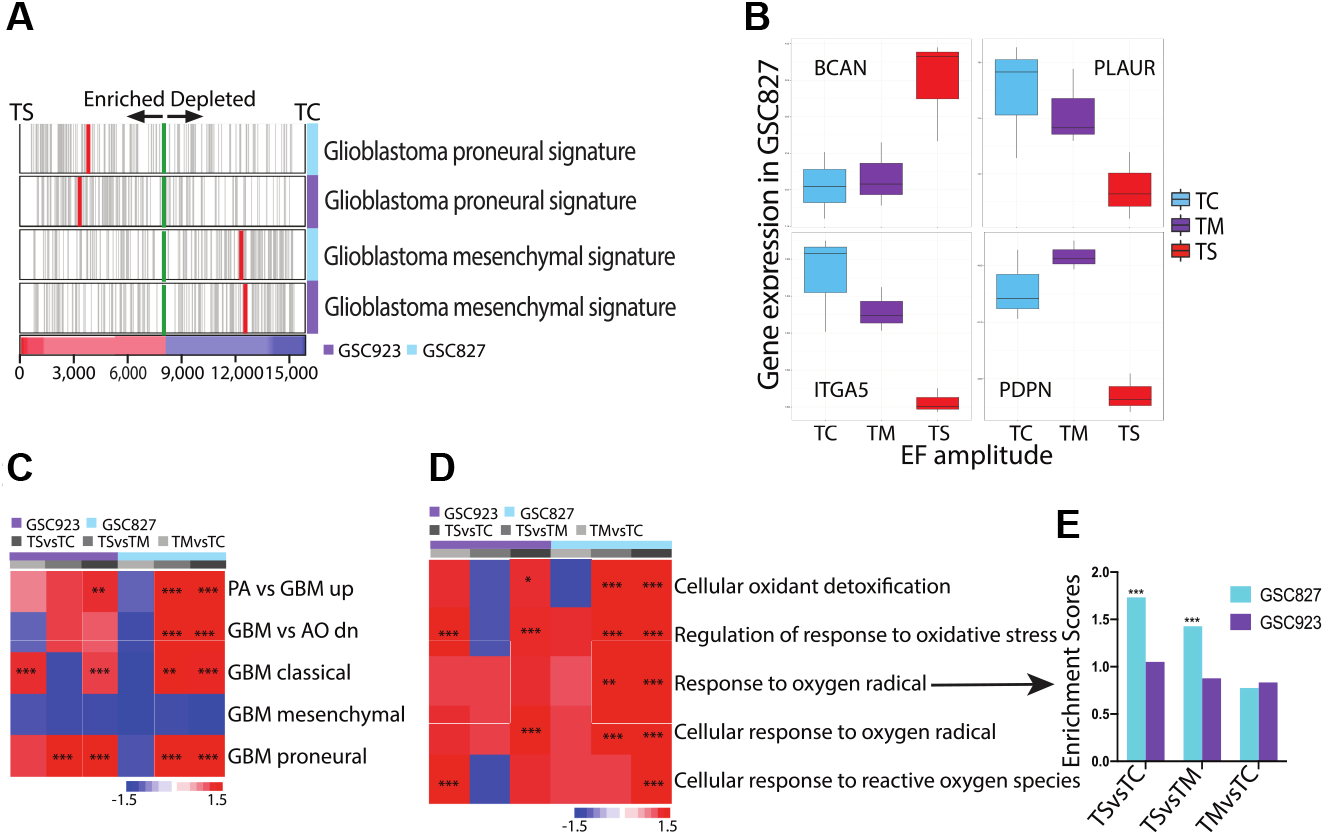
Treatment transformed the mesenchymal (MES) signature into proneural (PN) features in both cell lines and enhanced the cellular response to oxygen radicals in both cell lines. (A) Enrichment analysis of MES features and PN features. Green line indicates where expression correlation values change sign. Red line indicates where the leading-edge set starts. Image bar below indicates the sorted gene expression level. (B) Expression levels for select genes in the GBM subtype signature in GSC827. (C) Expression pattern of brain tumor related genesets. (D–E) Expression pattern and enrichment score (ES) of gene sets related to cellular response to oxygen radicals.

### Treatment increased the expression of genes related to cellular response to oxygen radicals in GSC827 and GSC923 cells

Treatment elevated the cellular response to oxidative stress in both cell lines. Five gene sets related to the cellular response to oxygen radicals were significantly enhanced for the TM vs. TC and TS vs. TM contrasts in GSC827 (Figs 5D–E). In GSC923, some, but not all of the same gene sets were significantly enhanced in the TM vs. TC and TS vs. TC contrasts (Fig 5D). This effect was more prominent in GSC827 than in GSC923 (Figs 5D–E), indicating a varied response to treatment across cell lines.

### Treatment increased the expression of genes related to cell cycle control and regulation in GSC827 cells

Our GBM network analysis derived two corresponding networks. For GSC827: GFAP, SOX2, TERT, MDM2, ATP1B2 and PROM1 were positively correlated while CD44, CDK4, CCND1, TP53, MTOR, and CLPTM1L were negatively correlated with the EF amplitude regions (Fig 6A). For GSC923, the positively correlated genes in the corresponding network were EFNB3, PDGFRA, IDH1, NOTCH1, FGF2, GFAP, ATP1B2 while the negatively correlated genes were CXCR4, CCND1, EPHA2, CLPTM1L (Fig S10E). The positively correlated genes for both cell lines were GFAP and ATP1B2 and the negatively correlated genes were CCND1 and CLPTM1L. GFAP is an astrocyte marker and ATP1B2 is a membrane protein responsible for establishing and maintaining the electrochemical gradients of Na^+^ and K^+^ ions across the membrane. CCND1 is a regulatory unit of CDK4 or CDK6 and is required for cell cycle G1/S transition. CLPTM1L is a transmembrane protein which is associated with poor prognosis, is overly expressed in ovarian serous adenocarcinoma, and confers resistance to chemotherapeutic killing [30].

**Figure 6.**
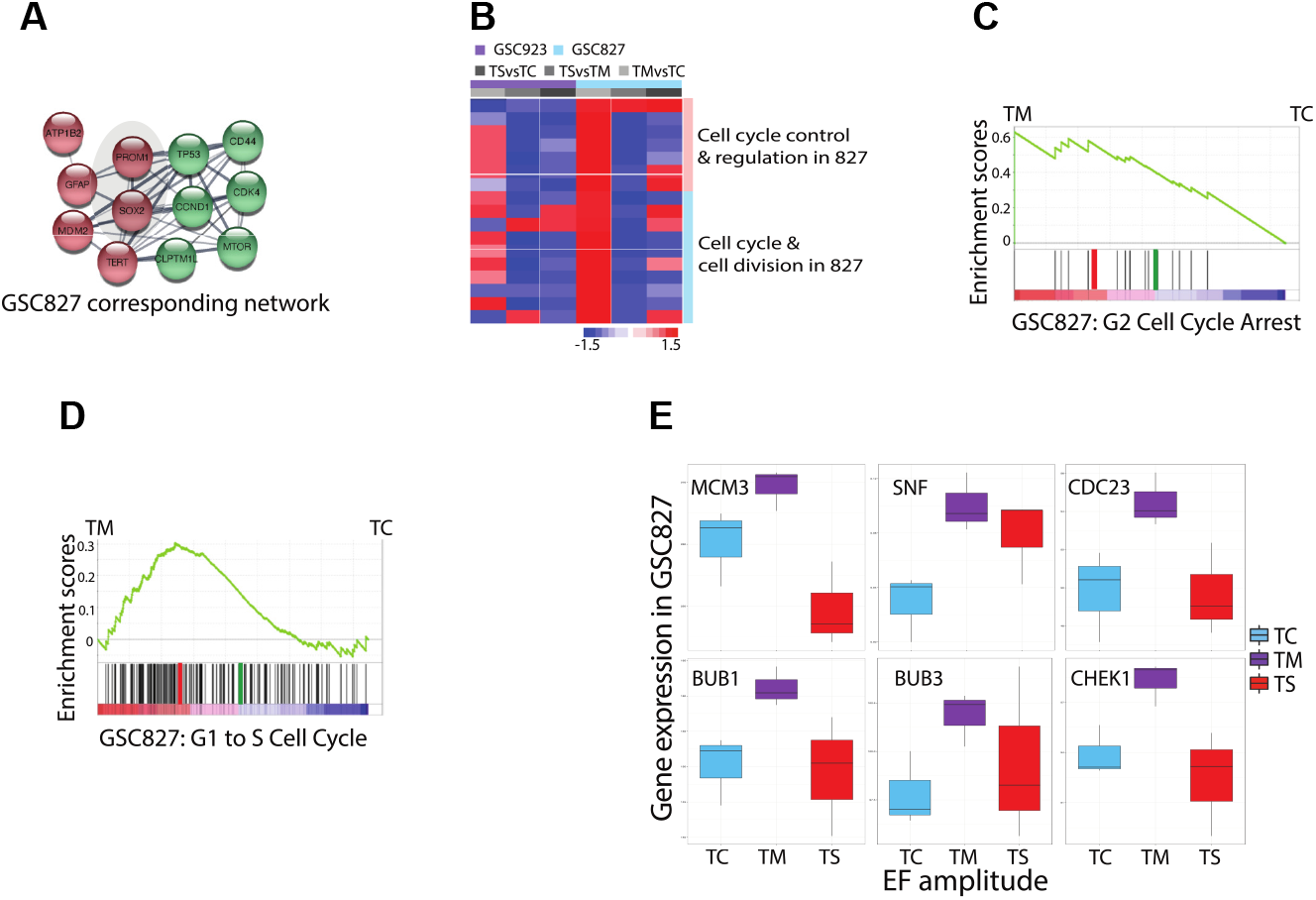
Treatment induced expression of genes related to cell cycle regulation in GSC827. (A) GBM corresponding network analysis for GSC827. Red nodes refer to upregulated genes and green nodes refer to down regulated genes. (B) Expression pattern of gene sets related to cell division and cell cycle control. (C) Enrichment analysis of the G2 cell cycle arrest gene set. Green line traces the running-sum of ES. (D) Enrichment analysis of the G1 to S cell cycle transition gene set. (E) Expression levels for select genes involved in cell cycle regulation in GSC827.

Our GSEA also identified seventeen gene sets associated with cell cycle regulation that were significantly enriched in GSC827 while contrasting TM vs. TC (Figs 6B–E). A full list of these gene sets is detailed in Table S1. Surprisingly, most of these gene sets were either down- or upregulated, but not reaching a significant level in the other two contrasts (TS vs. TM or TS vs. TC) within the same cell line (Fig 6B), indicating varied effects across EF amplitudes. In GSC923, a majority of these gene sets were not significantly enriched in the same direction as in GSC827 for the TM vs. TC contrast (Fig 6B). Expression levels for selected differentially expressed genes (DEGs) related to cell cycle control — CHEK1, BUB1, and BUB3 — are shown in Fig 6E. CHEK1 is a member of the Ser/Thr protein kinase family and is required for checkpoint mediated cell cycle arrest in response to DNA damage or the presence of replicated DNA. BUB1 and BUB3 encode for mitotic checkpoint Ser/Thr kinases that function by phosphorylating members of the mitotic checkpoint complex and activating the spindle checkpoint. All of these genes were significantly upregulated while contrasting TM vs. TC in GSC827 (Fig 6E). Expression levels for several other selected DEGs — SNF, MCM3, and CDC23 — are also shown (Fig 6E). SNF is a cell cycle checkpoint protein. CDC23 has an essential role in cell cycle progression through the G2/M transition. MCM3 is involved in the formation of replication forks and is essential for the initiation of DNA replication and cell cycle progression. In GSC923, none of these genes were significantly upregulated, again indicating a varied response to treatment across cell lines.

### Treatment increased the expression of genes related to NOTCH signaling and brain development in GSC923 cells

Treatment resulted in upregulated NOTCH and BMP signaling pathways in GSC923 (Figs 7A–E). Altogether, nineteen NOTCH signaling related pathways or gene sets from the MSigDB C2 and C5 databases were significantly enhanced in GSC923 across all TS vs. TC, TS vs. TM, and TM vs. TC contrasts (Figs 7A–B, S10F). These effects were less prominent in GSC827 and only four gene sets reached the significance level across all three contrasts (Fig S10F). Expression levels for selected upregulated DEGs involved in NOTCH signaling are shown in Fig 7E.

**Figure 7.**
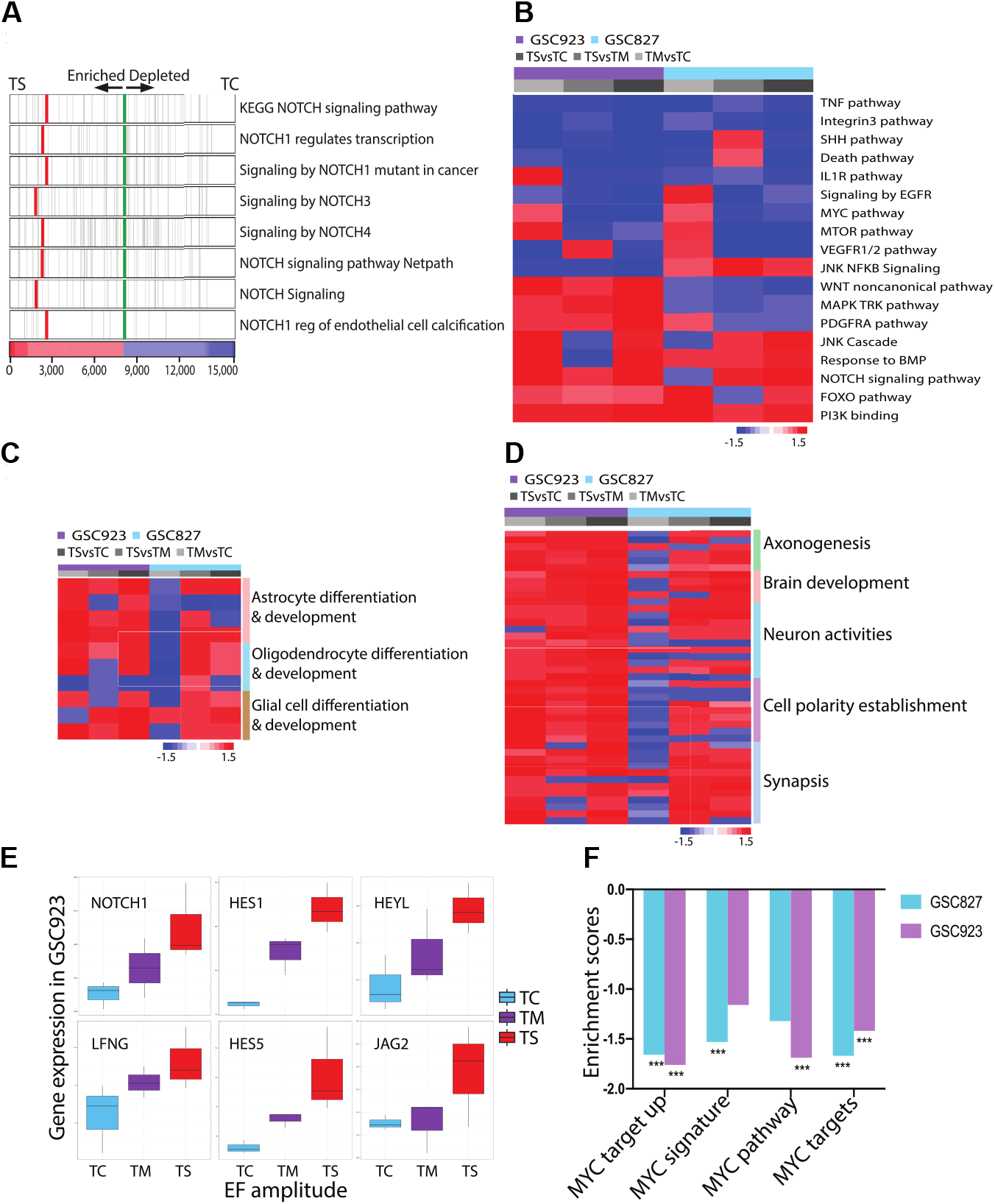
Genes related to NOTCH signaling, neuron activity, and brain development were promoted by treatment in GSC923. (A) Enrichment analysis of NOTCH signaling gene sets in the TS vs. TC contrast in GSC923. Green line indicates where the expression correlation values change sign. Red line indicates where the leading-edge set starts. Image bar below indicates the sorted gene expression level. (B) Significantly affected pathways by treatment across both cell lines. (C) Expression pattern of gene sets related to brain cell differentiation and development. (D) Expression pattern of gene sets related to neuron activities and brain development. (E) Expression levels for select genes in the NOTCH signalling gene sets in GSC923. (F) Enrichment scores (ES) for gene sets related to the MYC signalling pathway.

Gene sets related to brain cell differentiation and development, such as astrocyte and oligodendrocyte formation, were also enhanced by treatment in GSC923 (Figs 7C–D, S11A–B). Note that genes involved in NOTCH signaling, such as NOTCH1 and HES1/5, are in the edge gene list of brain cell differentiation function. The SOX1/6/9 transcription factors, which also contribute to brain development, were upregulated in GSC923 across contrasts (Fig S11B). Related genes such as OLIG1, GFAP, SOX2, and NKX2-2 were positively correlated with the EF amplitude region in GSC923 (Fig S11A). Genes related to cell adhesion, cell junction and synapsis, cell polarity establishment, axon extension and axonogenesis, neuron activities and the brain development functions were also enriched in GSC923 (Figs 4E, 7D). Two neuronal marker genes, MAPT — involved in the establishment and maintenance of neuronal polarity — and MAP2 — essential for neurogenesis — were positively correlated with the EF amplitude region (Fig S11A). Other genes related to neuronal activities, such as DCLK1, NLGN3, DISC1, EPHB1, SLIT2, and NTRK2, were also upregulated in GSC923 (Fig S11A).

### Other pathways and global functions affected by treatment

Overall, treatment with 200 kHz induced EMFs enhanced the expression of genes related to cellular polarity, synapsis, axonogenesis, neuron activities and brain development and downregulated the expression of genes related to functions of the ribosome, RNA activities, translation, abnormal glycosylation, and mesenchymal signatures (Fig 4E). Looking at more specific pathways (Fig 7B), the NOTCH signaling pathway, BMP signaling pathway, PI3K signaling pathway, and FOXO signaling pathways were enhanced to some degree in both cell lines. The PDGFRA signaling pathway, MAPK and TRK pathways, and WNT non-canonical pathways were enhanced in GSC923, but not in GSC827 (Fig 7B). JNK NFKB signaling was upregulated in GSC827, but not in GSC923 (Fig 7B). Pathways that were downregulated in both cell lines include the TNF pathway, integin3 pathway, SHH pathway, death pathway, and IL1R pathway (Fig 7B). The MYC pathway (Fig 7F), EGFR signaling, MTOR pathway, and VEGFR1/2 pathways were downregulated in the TS vs. TC and TS vs. TM contrasts, but not in the TM vs. TC contrast (Fig 7B). Expression levels of selected DEGs involved in these pathways — PDGFRA EGFR, MTOR, TERT, SHH, PIK3R1, FGF2, EPHA2, CXCR4, CD44 and IDH1 — are shown in Figs S12A–B.

Some other genes of interest were found to be differentially expressed. The major leading edge gene H19, which is a long, non-coding RNA that functions as a tumor suppressor [14], was upregulated in the TS condition for GSC827 (Fig S12C). The Na^+^-K^+^-2Cl^-^ transporter, NKCC1 (encoded by SLC12A2) was upregulated in the TM vs. TC and TS vs. TC contrasts in GSC827, but not in GSC923. NKCC1 regulates cell volume; NKCC1 is abundantly expressed in higher grade gliomas and its overexpression is thought to confer resistance to TMZ-induced apoptosis [25].

In summary, while several gene sets of interest were found to be up- or down-regulated, which may support certain mechanism(s) of action, the effects were not always consistent across cell lines, nor across contrasts (i.e., EF amplitudes).

## 1 Discussion

In the last 15 years, a growing body of work has suggested that TTFields (100 – 300 kHz, 1 – 3 V/cm) have an anti-proliferative effect, inhibiting the growth rate of human and rodent tumor cell lines [11, 18–20, 37]. This inhibitory effect was found to be frequency-dependent and to increase with EF amplitude. Novocure Ltd. translated these results toward various clinical applications, foremost in the treatment of GBM. Despite FDA approval of TTFields in combination with TMZ for the treatment of newly diagnosed GBM, adoption amongst neuro-oncologists remains low [37] due, perhaps, to a lack of understanding of the mechanism(s) of action of TTFields. Here, we have performed a study of alternating EF delivery via electromagnetic induction in the hopes of elucidating the *in vitro* effects of EFs/EMFs in this frequency and amplitude regime whilst mitigating possible thermal confounds endemic to previous EF delivery methods. Particular care was taken to eliminate thermal and osmolar artifacts.

The effects of 72 hours of uninterrupted 200 kHz EMFs with EF amplitudes spanning 0 – 6.5 V/cm were studied on cell cultures of primary rat astrocytes and the human glioma cell lines: U118, U87, GSC827, GSC923. Importantly, U87 and U118 were also used in the key pre-clinical studies that provided early evidence of the efficacy of TTFields. For the U87 cell line studied by Kirson *et al*. [20], we did not detect any anti-proliferative effect (Fig 2E). Surprisingly, we found that the apparent cell density increased by up to 30% within treated U87 cultures towards the higher EF amplitudes at the periphery of the dish (> 4 V/cm). For the U118 cell line also used by Kirson *et al*. [20], we found little to no effect on cell growth (≲ 5%). These results contradict previously published findings and do not support the claim that 200 kHz EFs exhibit single-modality efficacy in reducing the growth rate of these cell lines.

What could explain these contradictory results? Although a given EF should have the same biophysical effect(s) whether created capacitively (i.e., by time varying charge redistribution) or inductively (i.e., by time-varying magnetic fields), there are two main mechanistic differences between our method and the Inovitro™ EF delivery method: (1) In the Inovitro™ system, there is alternating stimulation between pairs of orthogonally-placed electrodes, and (2) the EMFs in our system are accompanied by a vertically-oriented, uniform, alternating magnetic field of about 0.6 millitesla.

These differences alone, however, are likely insufficient to explain the discrepancy in experimental results. According to Kirson *et al*. [18], alternating the EF orientation between two pairs of orthogonal electrodes increases the anti-proliferative effect of TTFields by just 20%, which is smaller than the discrepancies observed here: Kirson *et al*. [20] report growth rate inhibition of up to 50% in U87, whereas we report growth *enhancement* of up to 30%. Furthermore, these earlier studies reported increased efficacy at higher EF amplitudes. Thus, the higher amplitudes and continuous operation that were achieved here should, in theory, act to compensate somewhat for any loss in efficacy due to using a single alternating field orientation. With regards to the magnetic field, such a field is not predicted to have any direct effects on cells because (1) cells have a diamagnetic susceptibility nearly identical to that of water and (2) the 0.6 millitesla magnetic field amplitude is orders of magnitude smaller than those shown to have biological effects [50]. In addition, the magnetic field within the coil is predicted to be nearly uniform spatially, such that any effect of the magnetic field would be uniform throughout the dish and therefore incapable of explaining the observed effects, which vary radially.

Other reasons for the discrepancy in experimental results may arise from the direct contact between the electrodes and the dish in the Inovitro™ system, which, as discussed, complicates thermal and osmolar regulation and necessitates interrupted system operation in a refrigerated environment [33]. Critically, media evaporation is likely accelerated by the heating and the refrigerated environment. Novocure Ltd. advise that users of the Inovitro™ system compensate for this evaporation by changing media every 24 hours [33], which may cause osmolar shock to the cultures. Osmolar shock or stress alone could result in growth inhibition and apoptosis [6]. The oscillatory variations in temperature caused by the interrupted operation of the device may also contribute some anti-proliferative effect. Our system effectively mitigates these issues, although there remains some temperature variation within dishes (< 1 °C) and some additional osmolarity increase over 72 hours compared to controls (≈ 10%, see Figs 4 and 5 in [35]).

Moving to our other results, we found that for the human GBM stem cell lines GSC827 and GSC923, the apparent cell density decreased by up to 10 – 20% in the treated dishes at EF amplitudes exceeding 4 V/cm. For primary rat astrocytes, no effect on cell growth was observed. These results complement the U87 and U118 results by providing analogous findings for normal central nervous system cells (astrocytes) and for well-validated stem cell lines which better mimic the behavior of GBM *in vivo* [36].

We also assessed one candidate mechanism of action of TTFields involving interference with the multi-drug resistance of GBM cells — a feature conferred in part by ABC transporters. For GSC827 and GSC923 cells, the pumping activity of ABCG2 and ABCB1 was indirectly studied via the efflux of CellTracker and Hoechst over 2 hours. We found that there was little reduction in the apparent efflux of CellTracker and Hoechst in treated GSC827 and GSC923 cell cultures (Fig 3B), although marginally significant results observed at higher EF amplitudes may warrant further investigation.

The results presented here suggest strongly that different cells of different sources can respond to 200 kHz EMFs/EFs in various, even opposite ways with respect to cell growth. In our previous study [35], we studied human thyroid cells and observed a moderate reduction in cell density (< 10%) at low EF amplitudes (< 4 V/cm) and a greater reduction in cell density of up to 25% at higher amplitudes (4 – 6.5 V/cm), providing another example of a cell line with a distinct response profile. We are not the first to report such discrepant findings between cell lines. Similar results were presented by Neuhaus *et al*. [28], wherein TTFields were found to reduce the number of T98G cells but had the opposite effect on U251 cells, increasing their number (see Fig 9 in [28]). Our results also suggest that EF amplitudes which were previously thought to be effective *in vitro* (1 – 3 V/cm) may not be sufficient to elicit anti-proliferative effects when controlling for thermal and osmolar variations. The positive and negative growth rates observed at higher EF amplitudes (> 4 V/cm) in different cell lines merit further study.

To further explore possible mechanism(s) of action, RNA-Seq analyses were performed on cultures of GSC827 and GSC923 cells. We found several gene sets of interest to be up- or downregulated by treatment with 200 kHz induced EMFs. Our results indicate that treatment broadly modified the characteristics of both cell lines from mesenchymal (MES) enriched features to proneural (PN) enriched features (Figs 5A–B). In mice, MES glioma stem cells have a higher rate of proliferation and are more resistant to radiation than PN glioma stem cells [13]; mice injected with MES cells were found to develop brain tumors at a much faster rate. Among the four GBM molecular subtypes, the MES subtype is the most aggressive and strongly associated with poor prognosis compared to the PN subtype [24]. A transition from PN to the MES subtype can develop in patients following radiation or chemotherapy, and the tumor is often more resistant to treatment [38]. Therefore, the shift from MES to PN subtypes observed in our study is one potential mechanism of action for the reported clinical efficacy of TTFields independent of any direct reduction in cell growth. Another effect found in both cell lines, but only significantly in GSC827, was an increased expression of genes related to the cellular response to oxygen radicals. While reactive oxygen species are known to be involved in pro-tumorigenic and anti-tumorigenic signalling [27, 34], it is unclear, at present, how and whether this relates to the effects of TTFields in the clinic.

Other effects on gene expression were found only in one cell line or only at some EF amplitudes. We found that genes related to cell cycle control and regulation were upregulated in GSC827, but not in GSC923 cells (Fig 6). Furthermore, the effect was only present at intermediate EF amplitudes (TM, 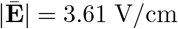) and disappears at higher EF amplitudes (TS, 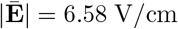) within the GSC827 cell line. Effects on the mitotic process and cell cycle are promising candidates for the possible mechanism of action(s) of TTFields and were originally proposed as such [18, 20]. Thus, it is surprising that our RNA-Seq analyses did not find a consistent modulation of gene expression for genes related to cell cycle regulation across cell lines, nor across EF amplitudes. Gene sets related to anaplastic oligodnedroglioma and pliocytic astrocytma signatures — which are lower grade tumors compared to GBM — were found to be upregulated in GSC827, but not significantly in GSC923 (Fig 5C).

Genes related to NOTCH signalling and brain cell development were found to be upregulated in GSC923 but not to the same degree in GSC827 (Fig 7). NOTCH signaling is fundamental for maintaining neural stem cells and for regulating cell fate specification, proliferation, and apoptosis [21, 31]. The NOTCH pathway has been recognized as having a key role in the early stages of neurogenesis and plays a role in tumor suppression [26, 29, 31, 44]. NOTCH1, in particular, is essential for neuron and glia formation and facilitates the onset of neurogenesis by offering a signal of astroglia differentiation independent of ciliary neurotrophic factor [26, 44]. Consistent with these reports, our GSEA revealed strong enriched signatures of both NOTCH signalling and neuroglial development, manifesting as astrocyte and oligodendrocyte cell differentiation and brain development (Fig 7) in GSC923. In a subset of GBM patients, high expression of specific NOTCH target genes is associated with a better prognosis. In a glioma mouse model, NOTCH2 overexpression inhibited glioma formation [8]. A recent CRISPR screening identified NOTCH1 as a tumor suppressor in GBM [5]. Taking the above findings into account, enhanced NOTCH signaling may represent a novel route of EF effects, which could benefit patients by restoring cell capacity via differentiation. Again, however, this effect was more prominent in GSC923 than GSC827.

In summary, the RNA-Seq analyses carried out here revealed several new potential mechanisms of action that may lead to improved prognosis without, necessarily, an accompanying reduction in cell growth, which was observed only to a moderate degree at higher EF amplitudes in our imaging studies of GSC827 and GSC923 cells (Fig 2E). In particular, the transition from MES enriched features to PN enriched features and an increased response to oxygen radicals was found in both cell lines and the direction of effect was consistent with increasing EF amplitude. Other effects, such as enhanced NOTCH signalling, neural differentiation, or cell cycle regulation, were also observed, but were not consistent across cell lines nor across EF amplitudes.

## 2 Conclusions

The response to treatment with 200 kHz induced EMFs, in terms of cell growth and more general gene expression, was found to be highly variable and specific both to cell line and EF amplitude when temperature and osmolarity were carefully controlled. We did not find that cell growth was significantly inhibited in the EF amplitude regime of 1 – 3 V/cm across GBM cell lines, contradicting key prior findings. Indeed, at higher EF amplitudes, U87 cells responded to treatment with *enhanced* cell growth. However, we cannot rule out that effects which do not result in an observable reduction in cell growth could be caused by such EMFs. We report preliminary RNA-Seq analyses on human glioma stem cell lines which reveal some effects along these lines. For example, a transition from mesenchymal to proneural cell signatures was observed in both the GSC827 and GSC923 GBM stem cell lines. Overall, the varied responses to treatment suggest the need to redouble our efforts to understand any possible effects of alternating electric fields on tissues and cells in this frequency regime, and to begin assessing such effects on a per cell line basis, and in other biological and clinical contexts.

## Contributions

Conceptualization, R.R., P.J.B. and M.R.G; Methodology, R.R., T.X.C., A.L., R.H.P., M.G-C, T.P.; Software, T.X.C and A.L.; Validation, R.R., T.X.C and A.L.; Formal analysis, R.R., T.X.C. and A.L.; Investigation, R.R. and N.B.; Resources, R.R., R.H.P., Z.Z., H.W. and J.C.; Data curation, R.R., T.X.C. and A.L..; Writing—original draft preparation, R.R., T.X.C and A.L.; Writing—review and editing, R.R., T.X.C., A.L., N.Y.M., N.H.W., P.J.B. and M.R.G.; Visualization, R.R., T.X.C., A.L., M.G-C; Supervision, P.J.B.; Project administration, M.R.G and P.J.B.; Funding acquisition, P.J.B. All authors have read and agreed to the published version of the manuscript.

## Funding

This work was supported by the IRP of the NICHD and the NCI.

## Data Availability

Data available upon reasonable request.

## Acknowledgements

We would like to thank Dr. Phillip R. Lee and Dr. R. Douglas Fields, who provided astrocyte expertise and contributed to the discussion. We would also like to thank Raisa Freidlin, D.Sc. who was engaged in early image processing activities.

## Conflicts of Interest

The authors declare no conflict of interest.

## Notes

### Competing Interest Statement

The authors have declared no competing interest.

