## Supplementary Material for "“Tumor Treating Fields” delivered via electromagnetic induction have varied effects across glioma cell lines and electric field amplitudes"

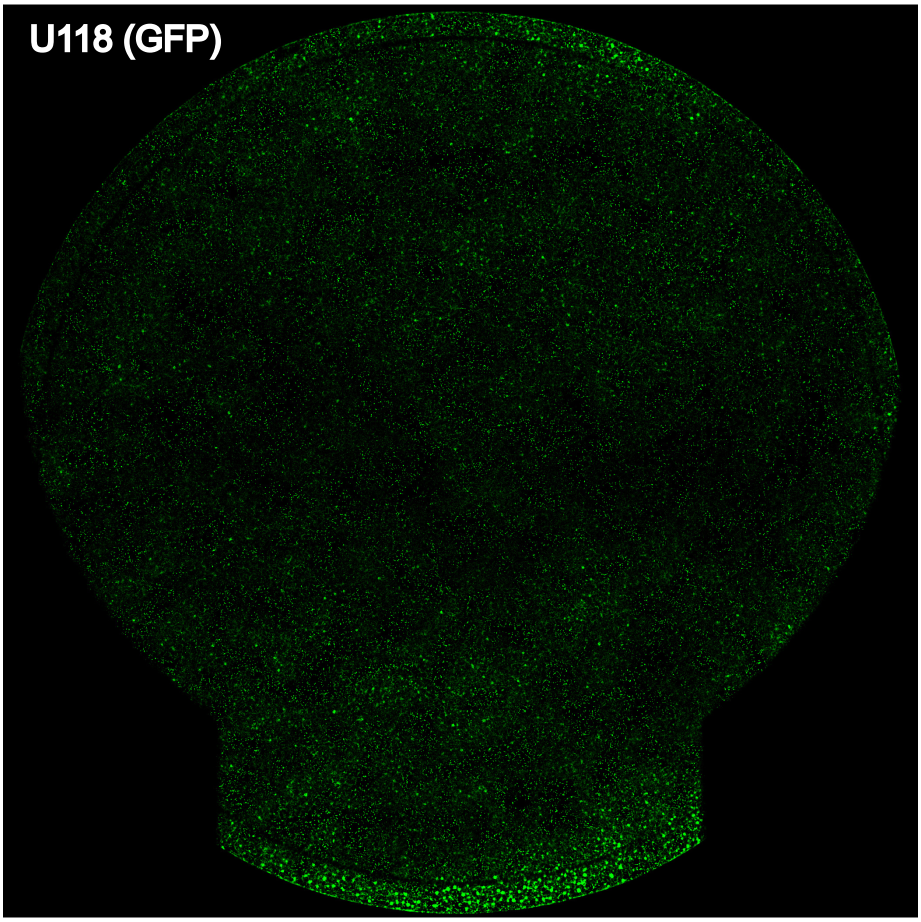


Figure S1: Exemplar image of a whole dish from a U118 (GFP) control culture. Note that the fitments occlude the bottom portion of the dish and that a rim from the manufacture of the dish occludes an outer rim. Exclusion of certain bands as shown in Fig. 2D is justified due to these dish characteristics.

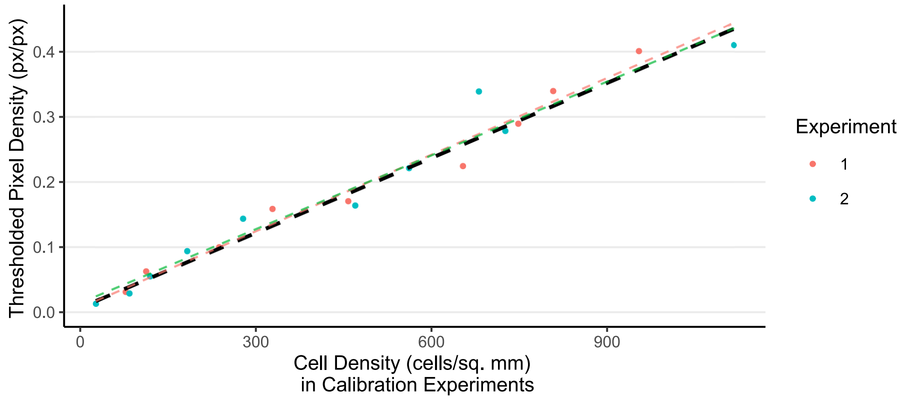


Figure S2: Calibration of U87 cell counts compared to segmented px from GFP expression. In separate calibration experiments, different initial seeding densities ranging from $1\times10^4$ to $1\times10^5$ cells per dish were used. Cells were detached with Accutase, incubated for 24 h, and then incubated for an additional 72 h. At the end of 72 h, the cell culture dishes were scanned using confocal imaging as described in the main text and all cells were subsequently lifted and counted using a Countess™ II FL Automated Cell Counter (Thermo Fisher). Final cell densities ranged from $< 100$ to 1200 cells/sq. mm. Two series of calibration experiments were performed for a total of 38 cell counts and images. For image processing, the same pipeline as in the main text was used but with fixed intensity thresholds of 0.06 and 0.51 (i.e., keeping px between the thresholds) and without binning. Combining the two experiments, a linear fit (black dashed line) between cell density and thresholded px density was obtained: $[\text{Px Density}] = 3.84 \times 10^{-4} \cdot [\text{Cell Density}] + 1.1 \times 10^{-2}$, Adj. $R^2 = 0.96$. Individual fits to each set of experiments are also shown.

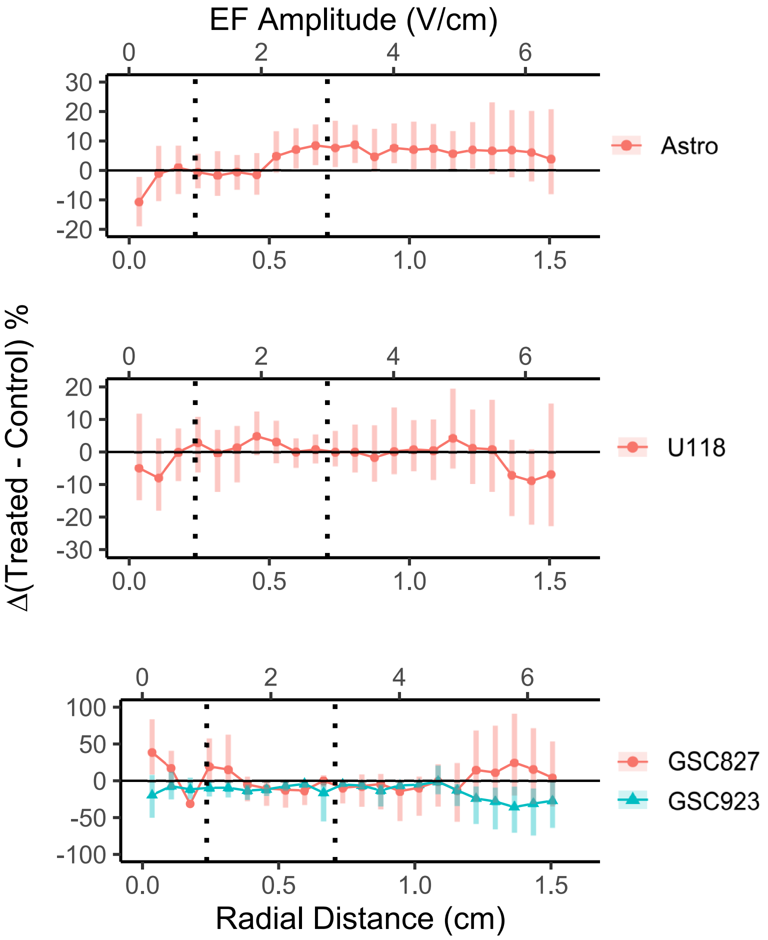


Figure S3: Cell density changes measured using segmented px densities obtained from Hochest staining. The same imaging pipeline was performed as described in the main text. Note that the effect observed for GSC827 in the main text disappears. Hoechst staining was not performed for U87.

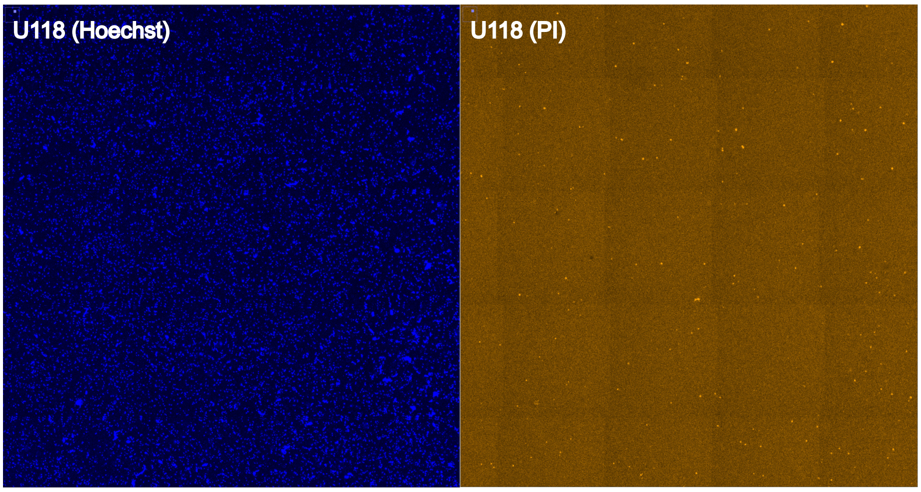


Figure S4: Exemplar ROIs from Hoechst and propidium iodide (PI) staining in a treated U118 cell culture. Note the sparsity of PI staining compared to Hoechst. PI (V13241; Thermo Fisher Scientific) staining was performed after GFP imaging. At the end of experiments and 5 min before microscope imaging, 750 $\mu$L of DMEM was removed from the dishes and mixed with 0.5 $\mu$L of 1 mg/mL stock solution and added back to the dish to a final concentration of $2 \times 10^{-4}$ mg/mL. PI staining was performed in $N = 12, 9$ pairs of U118 and U87 cultures, respectively. In all such cases, few to no dead cells were found, and further statistical analyses were not carried out.

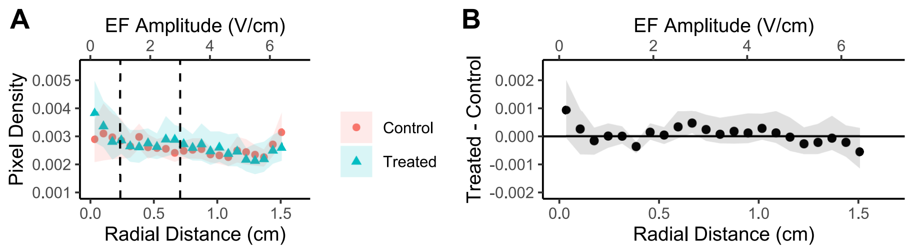


Figure S5: Results of Annexin V staining in U118 cell cultures. Alexa Fluor™ 594 (A13203 Invitrogen) staining was performed with 10 $\mu L$ Annexin V in 200 $\mu L$ of DMEM for 10 minutes — washing 2× with 750 $\mu L$ of DMEM before imaging. Four experiments were performed. One experiment was excluded due to high amounts of noise in the dish periphery for a total of 9 dish-to-dish comparisons. (A) Segmented px density was calculated using the same image processing pipeline as described in the main text, except that area opening was performed for a smaller size of 2 px. Error bars = 90% confidence intervals from 1000 re-samplings. Px density values were not normalized for this analysis. (B) Difference between treated and control cultures in terms of raw px density. Error bars = 90% confidence intervals from bootstrapped pair-wise differences with 1000 re-samplings.

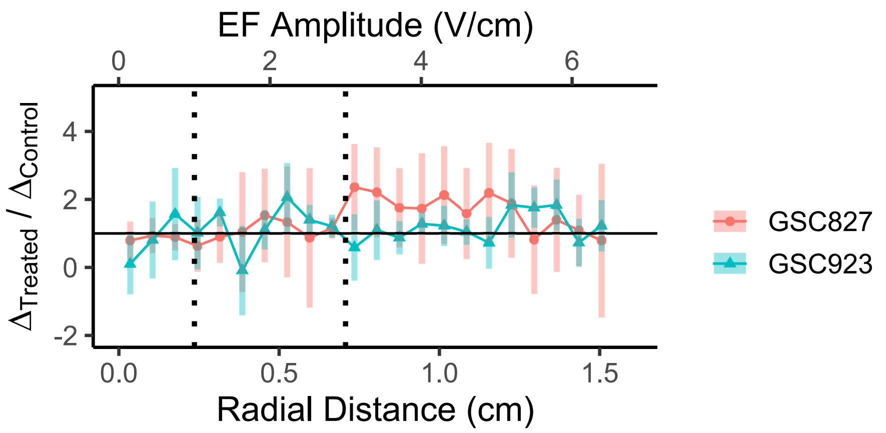


Figure S6: Analysis of Hoechst pumping as described in the main text. Outliers with $\Delta_{\text{Treated}}/\Delta_{\text{Control}}$ ratios exceeding an absolute value of 10 (arising due to the small values of $\Delta$) were excluded for a total exclusion of 31/638 data points ($\approx$ 5%). No difference in apparent pumping activity is observed for either cell line.

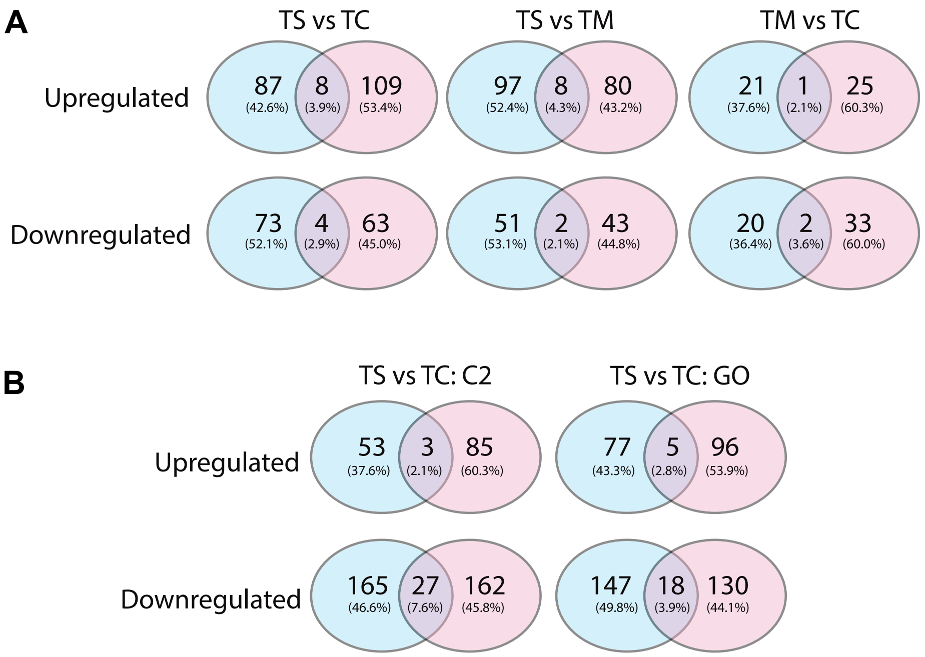


Figure S7: The two glioma stem cell lines, GSC827 and GSC923, are transcriptomically different. (A) Venn diagrams of up/down regulated genes in all three contrasts: TS vs. TC, TS vs. TM, and TM vs. TC. Cyan represents differentially expressed genes in GSC827 and pink represents GSC923. (B) Venn diagram of up/down regulated gene sets in GSEA using MSigDB C2 and C5 databases.

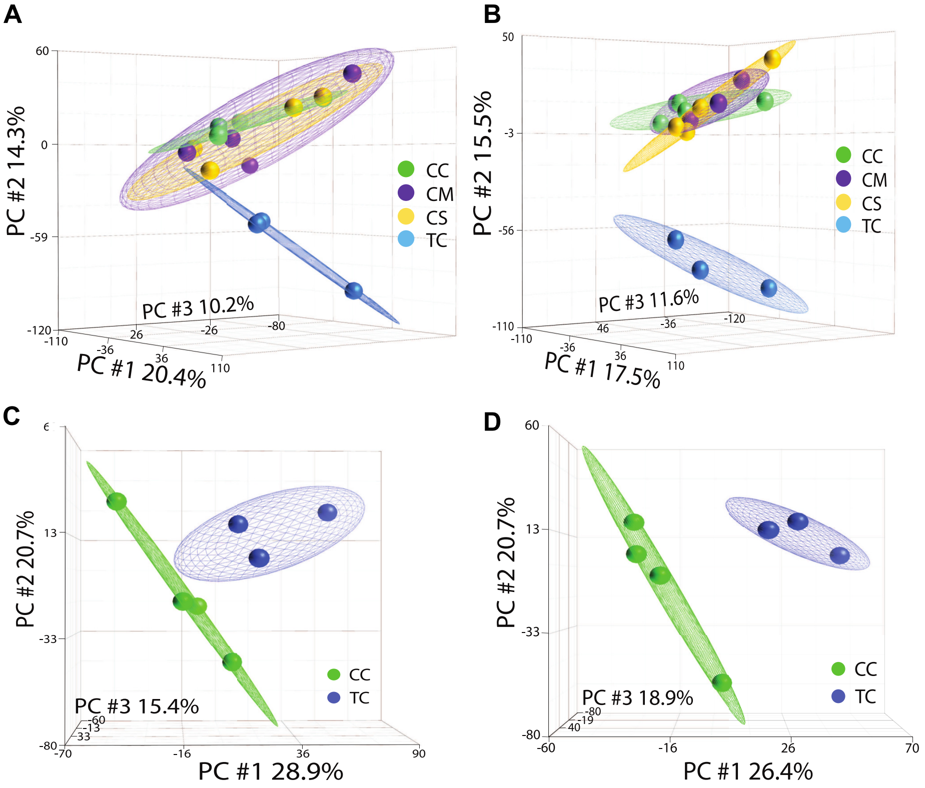


Figure S8: Differences between treated center (TC) and control dish regions (CC, CM, CS). (A, C) Unsupervised PCA of GSC827. (B, D) Unsupervised PCA of GSC923. Note the difference between TC and CC in both cell lines.

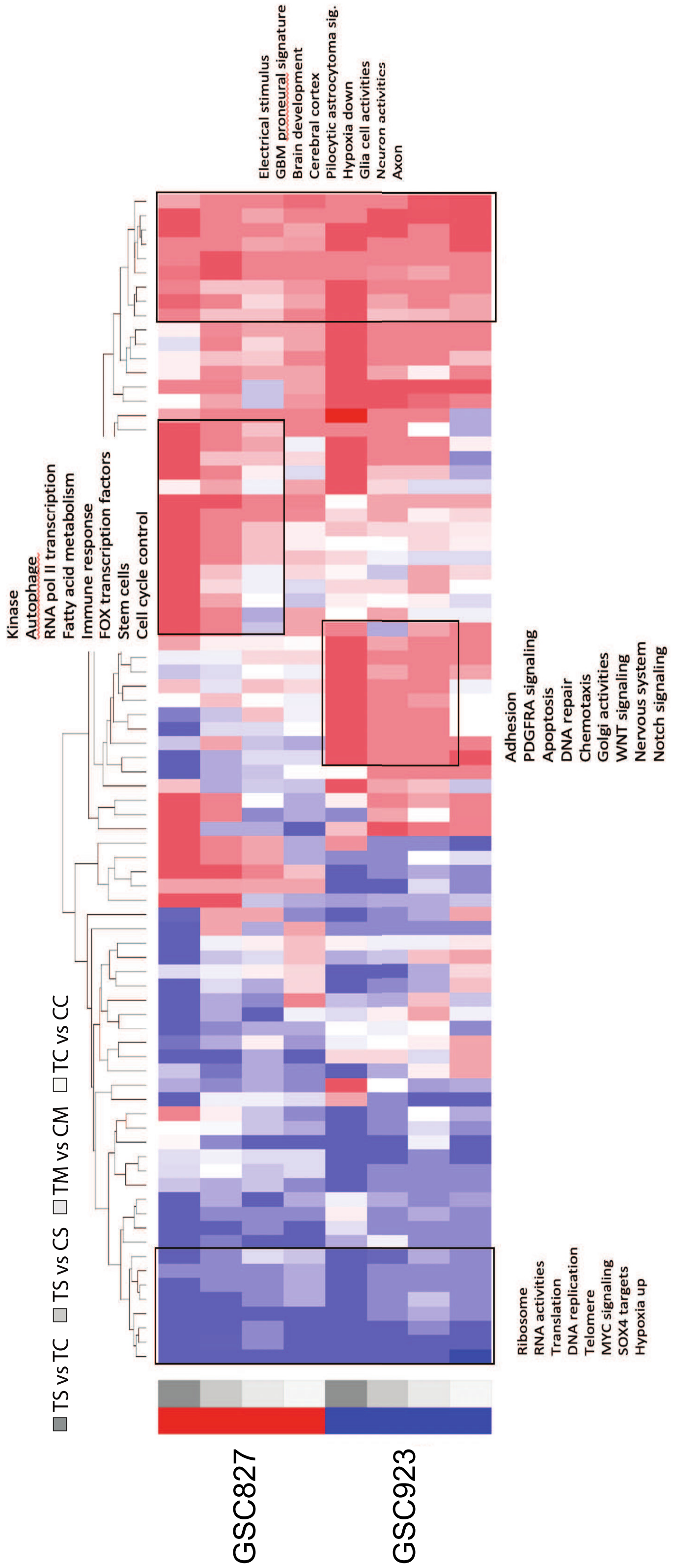


Figure S9: Functional summary of TS vs. TC, TS vs. CS, TM vs. CM and TC vs. CC in both cell lines, GSC827 and GSC923.

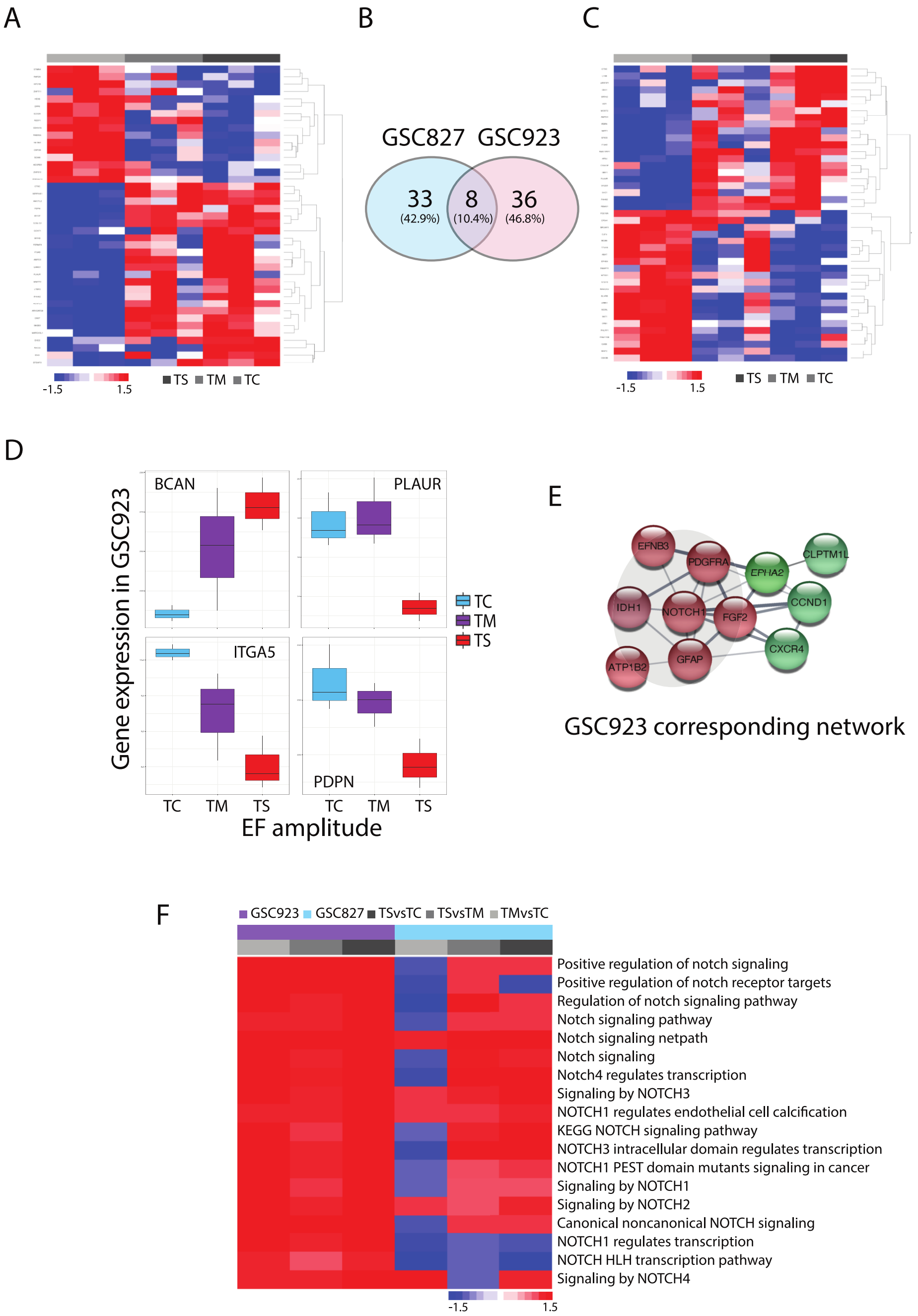


Figure S10: The expression of mesenchymal and proneural signatures and NOTCH signaling gene set enrichments. (A–C) PN and MES signature affected by EF amplitude in GSC827 (left) and GSC923 (right). (D) Selected gene expression of PN and MES signature in GSC923. (E) Corresponding network analysis for GSC923. Red nodes refer to upregulated genes and green nodes refer to downregulated genes. (F) Enriched expression of NOTCH signaling gene sets in GSC827 and GSC923.

Table S1: Cell cycle gene sets affected by EF amplitude in GSC827

| Gene set | TS vs. TC | $p$-val | TS vs. TM | $p$-val | TM vs. TC | $p$-val |
|---|---|---|---|---|---|---|
| POSITIVE_REGULATION_OF_CELL_CYCLE_ARREST | 1.42 | $< 0.01$ | 1.15 | 0.19 | 1.40 | $< 0.01$ |
| RTM_CELL_CYCLE_CHECKPOINTS | -0.82 | 0.82 | -1.30 | $< 0.01$ | 1.57 | $< 0.01$ |
| RTM_G2_M_CHECKPOINTS | -0.89 | 0.63 | -1.41 | $< 0.01$ | 1.60 | $< 0.01$ |
| RTM_G1_S_DNA_DAMAGE_CHECKPOINTS | 1.11 | 0.26 | -1.11 | 0.35 | 1.75 | $< 0.01$ |
| TP53_REGULATES_GENE_TRANSCRIPTION_IN_G2 _CELL_CYCLE_ARREST | 1.20 | 0.34 | -1.10 | 0.30 | 1.95 | $< 0.01$ |
| KEGG_CELL_CYCLE | -0.97 | 0.62 | -1.49 | $< 0.01$ | 1.32 | 0.05 |
| FISCHER_G1_S_CELL_CYCLE | 1.17 | 0.10 | -0.98 | 0.31 | 1.36 | $< 0.01$ |
| RTM_MEIOSIS | 0.76 | 0.77 | -1.28 | 0.11 | 1.41 | $< 0.01$ |
| RTM_S_PHASE | -1.03 | 0.38 | -1.37 | $< 0.01$ | 1.48 | $< 0.01$ |
| RTM_MITOTIC_METAPHASE_AND_ANAPHASE | -0.10 | 0.53 | -1.38 | $< 0.01$ | 1.51 | $< 0.01$ |
| RTM_CYCLIN_A_CDK2_ASSOCIATED_EVENTS_AT_S _PHASE_ENTRY | 0.46 | 0.90 | -1.089 | 0.24 | 1.53 | $< 0.01$ |
| RTM_SEPARATION_OF_SISTER_CHROMATIDS | -0.91 | 0.74 | -1.37 | $< 0.01$ | 1.57 | $< 0.01$ |
| WHITFIELD_CELL_CYCLE_LITERATURE | -0.56 | 0.74 | -1.00 | 0.40 | 1.72 | $< 0.01$ |
| GOCC_ANAPHASE_PROMOTING_COMPLEX | -0.68 | 0.91 | -1.05 | 0.27 | 1.42 | $< 0.01$ |
| RTM_MEIOTIC_RECOMBINATION | 1.10 | 0.3 | -1.06 | 0.39 | 1.76 | $< 0.01$ |
| RTM_GTSE1_IN_G2_M_PROGRESSION_AFTER_G2 _CHECKPOINT | -0.84 | 0.70 | -1.27 | 0.31 | 1.61 | $< 0.01$ |
| RTM_APC_C_MEDIATED_DEGRADATION_OF_CELL _CYCLE_PROTEINS | -0.60 | 0.81 | -1.24 | 0.21 | 1.68 | $< 0.01$ |

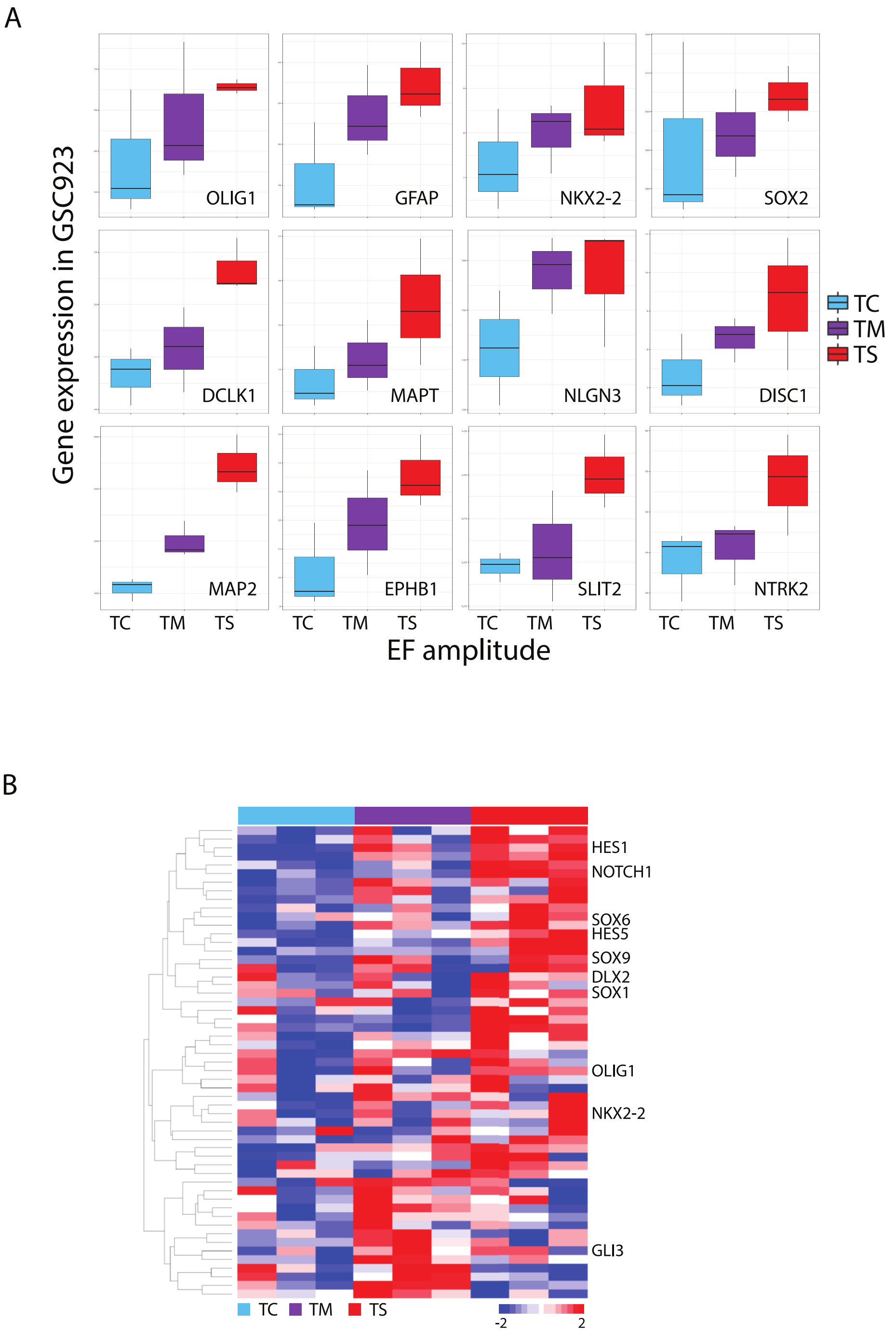


Figure S11: Selected gene expression related to brain cell differentiation, neuron activities and brain development. (A) Selected gene expression involved in neuron activities, and brain development that are positively correlated with the EF amplitude (i.e., the dish regions: TS > TM > TC) in GSC923. (B) Supervised hierarchical clustering of genes involved in brain cell differentiation in GSC923.

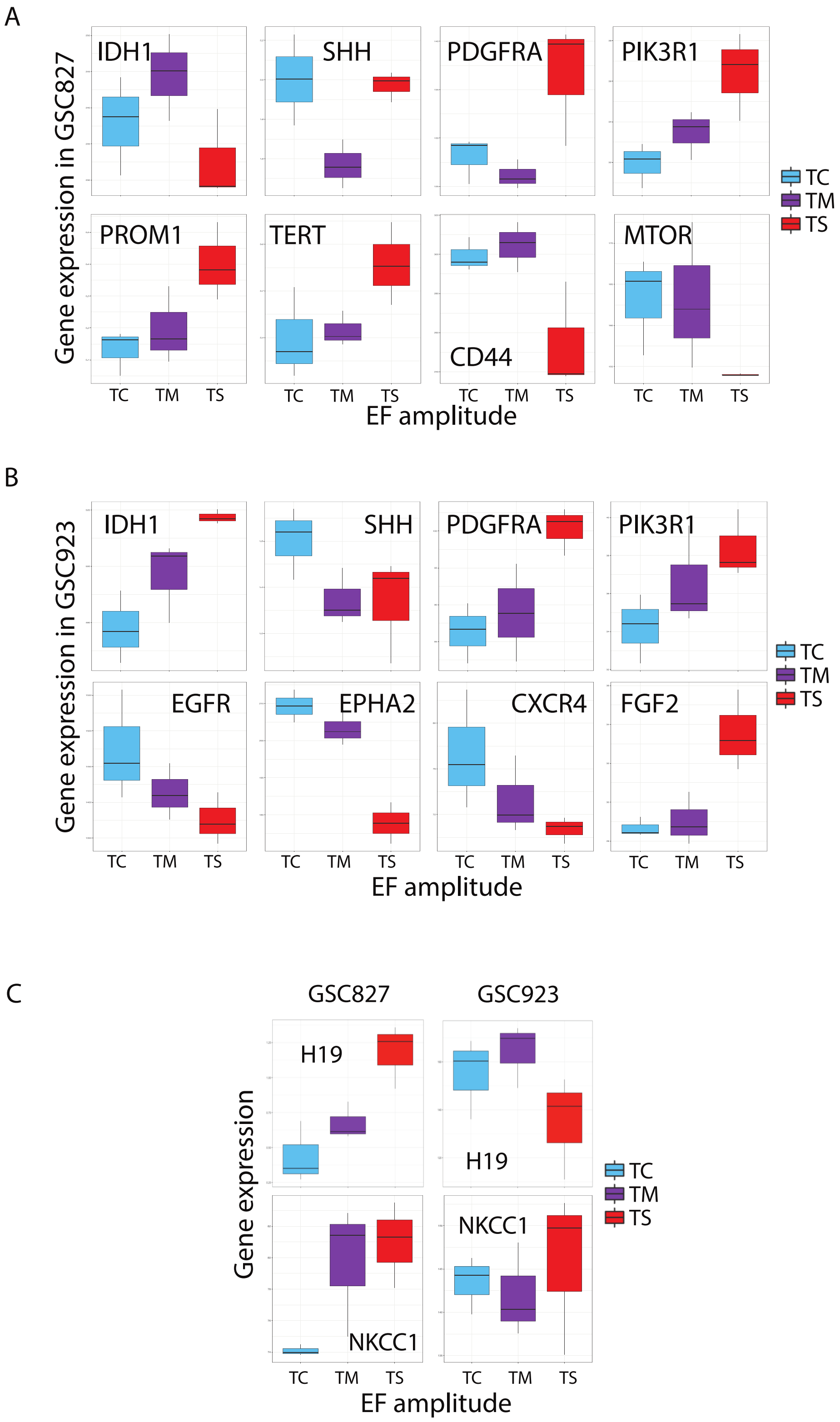


Figure S12: Selected gene expression in both cell lines. (A) Selected gene expression in GSC827. (B) Selected gene expression in GSC923. (C) H19 and NKCC1 expression in both cell lines.
